# Brassinosteroids mediate proper coordination of sepal elongation

**DOI:** 10.1101/2025.07.13.664398

**Authors:** Byron Rusnak, Lilijana Oliver, Kyle Procopio, Adrienne Roeder

## Abstract

*Arabidopsis* sepals must grow in a coordinated and robust fashion to a consistent size and shape to close and protect the developing flower bud. To understand how this robust coordination occurs, we use the loss of robustness mutant *development related myb-like1* (*drmy1*), which exhibits variable sepal initiation and growth, causing failure of the sepals to close the flower bud. Specifically, *drmy1* has overgrown outer (abaxial) sepals and undergrown inner (adaxial) sepals, leading to a large discrepancy in the sizes of different sepals within individual flower buds. Using single cell and spatial RNA-seq, we found changes in expression of key genes related to brassinosteroid (BR) signaling in *drmy1*, particularly in cell types important to young flower bud development such as epidermal cells, boundary cells, and meristematic cells. Confocal imaging of a BRI1-EMS-SUPPRESSOR1 (BES1) ratiometric reporter confirms that BR signaling is upregulated and more variable in young *drmy1* sepals. Subsequently, we found that altering BR signaling in *drmy1* by crossing with BR mutants or adding brassinolide (a potent brassinosteroid) or brassinazole (a brassinosteroid biosynthesis inhibitor) can partially rescue this elongation defect by differentially altering the relative growth of the inner and outer sepals. Increasing BR signaling rescues by increasing the growth of the inner sepal but not the outer sepal, while decreasing BR signaling rescues by decreasing the growth of the outer sepal but not the inner sepal. These results suggest that brassinosteroids mediate the robust coordination of the growth rates between inner and outer sepals during early development, ensuring proper flower bud closure.

## Introduction

In biology, robustness is the concept that living organisms can produce structures of consistent size and shape within and between individuals despite cell heterogeneity, gene expression stochasticity, and perturbations from the external environment (Lachowiec et al. 2016). A common theme for generating robust forms is the coordination of growth between various parts of a structure (Kong et al. 2024a). Coordination operates at multiple scales. For instance, within a single tissue layer such as the leaf or the sepal epidermis, cell expansion and divisions are coordinated to achieve proper size and flatness (Harline et al. 2023; Horiguchi et al. 2005; Hong et al. 2016; Burda et al. 2024). Additionally, tissues such as the L1, L2, and L3 layers of the meristem coordinate cell division planes and growth between layers through a combination of hormone and peptide signaling (Savaldi-Goldstein and Chory 2008; Schoof et al. 2000). Growth of the adaxial and abaxial epidermal layers of sepals must also be coordinated to achieve proper curvature (Xu et al. 2024; Yadav et al. 2024). Coordination is also necessary between separate structures on an individual, such as the wings on *Drosophila melanogaster* which must carefully coordinate equally but mirrored growth to achieve symmetric flight (Parker and Struhl 2020; Crickmore and Mann 2006).

Sepals provide an ideal model for understanding robustness and coordination, as proper coordination of cell expansion within a sepal is necessary to achieve the correct size and shape (Hervieux et al. 2016; Hong et al. 2016; Burda et al. 2024; Roeder 2021). There must also be proper coordination of initiation and growth between different sepals in an individual flower to enclose the flower bud properly (Zhu et al. 2020; Kong et al. 2024b; Kong et al. 2024c).

Previously, we characterized the *development-related myb-like 1* (*drmy1*) mutant, which produces sepals of variable size and shape, preventing proper closure of the developing flower bud (Figure 1A) (Zhu et al. 2020; Kong et al. 2024b; Kong et al. 2024c). We showed that delayed and variable sepal initiation contributes to this phenotype (Zhu et al. 2020). Bulk RNA sequencing (RNA-seq) identified a reduction in translation and an increase in the expression of the boundary gene *CUC1* in *drmy1* (Kong et al. 2024b). Follow-up work using fluorescent reporters for auxin signaling (DR5) and cytokinin signaling (TCS) revealed that impaired translation leads to elevated cytokinin and reduced auxin signaling, driving sepal initiation delays and variability (Kong et al. 2024b). However, *drmy1* defects in post-initiation sepal elongation remain understudied. Additionally, these changes in hormone signaling escaped detection in bulk RNA-seq experiments, likely because cell-type specific signatures were obscured by the mixed cell population. These limitations can be overcome with single cell RNA-seq (scRNA-seq) and spatial transcriptomics, which preserve cell type and spatial context, respectively, and are important tools in developmental biology (Rusnak et al. 2024).

**Figure 1:**
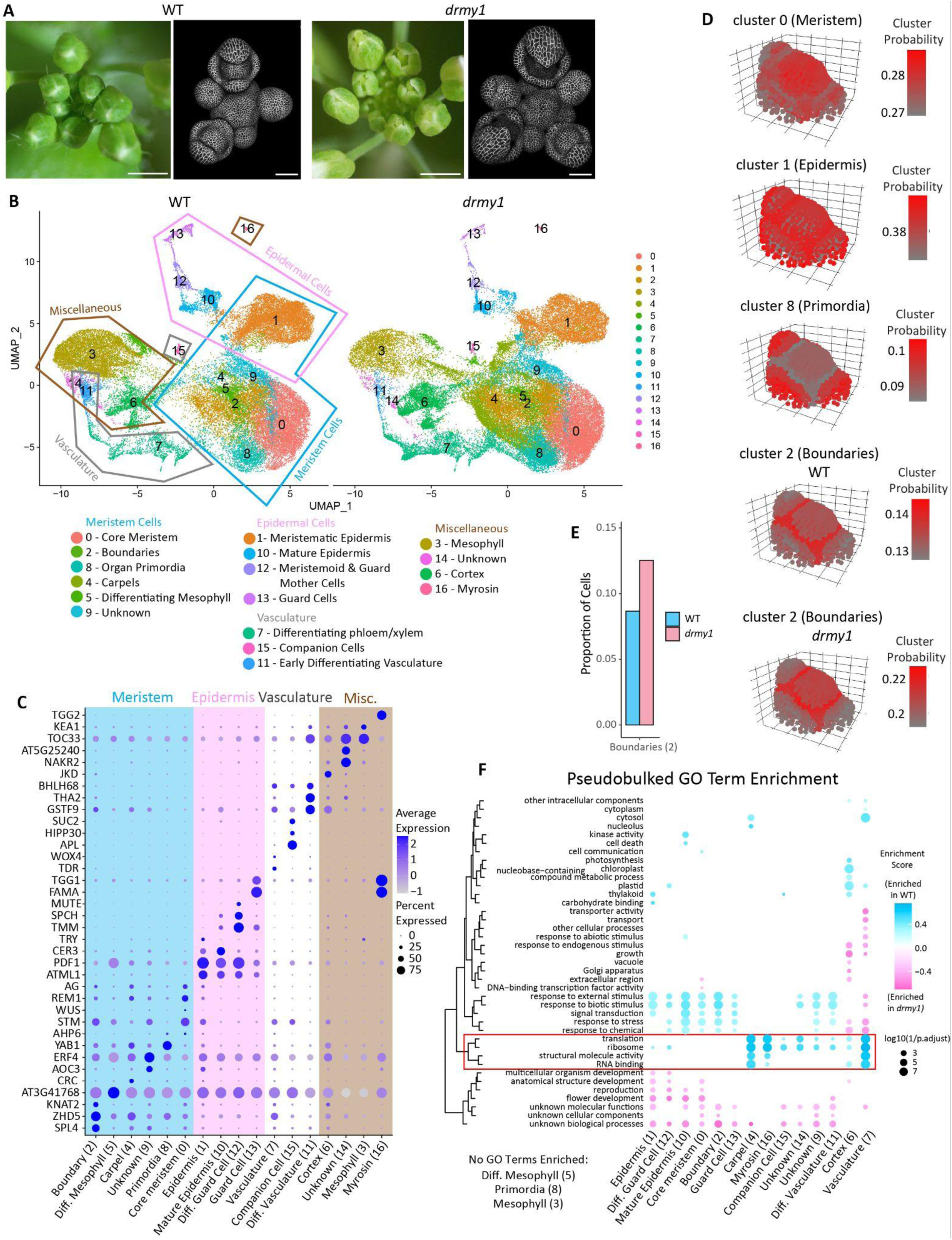
Single cell RNAseq shows *drmy1* has more boundary cell identity and reduced translation. A) Wild type (WT) and *drmy1* inflorescences showing flower buds at stages 11-12 (stereomicroscope images) and at stages 0-5 (volume renderings of confocal images). Confocal images were taken of plants expressing the plasma membrane marker *35S::mcitrine-RCI2A*. Note the defects in sepal sizes that prevent stage 11-12 buds from closing and the elongated abaxial (outer) sepals in stage 4-5 buds. Scale bars 1 mm (stereomicroscope images) or 50 μm (volume renderings of confocal images). B) UMAP projection of single cell RNAseq (scRNAseq) data with cluster identities labeled. C) Dotplot of marker genes used to identify cluster identities using SCT normalized expression values. D) Single cell clusters that mapped with high probability onto the floral meristem atlas. Note the higher probabilities for which the boundary cluster maps to the atlas in *drmy1*. All other mappings shown are WT. E) Proportion of cells assigned to the boundary cluster in WT (0.087) and *drmy1* (0.125). F) Gene Ontology Term enrichment of differentially expressed genes in *drmy1*. Blue indicates higher enrichment in WT, pink indicates higher enrichment in *drmy1*. (Supplementary Data 1) See also: Figure S1

In this paper, we applied comparative single cell RNA-seq (scRNA-seq) and Visium spatial RNA-seq to wild type (WT) and *drmy1* flower buds during early sepal development to identify cell type and location specific transcriptomic changes. We confirm subtle shifts in auxin and cytokinin signaling that previously escaped detection by bulk RNA seq, reveal novel miscoordination in brassinosteroid (BR) signaling pathways, and validate these findings with a BR responsive ratiometric fluorescent reporter. Further, we show that genetic and chemical modulation of BR signaling differentially affects inner versus outer sepal elongation within the same flower bud. Increasing BR signaling partially rescues sepal elongation defects by preferentially increasing elongation of the inner sepal, while decreasing BR signaling partially rescues by decreasing overgrowth of the outer sepal. These results highlight a novel role for brassinosteroids in coordinating the growth between sepals in the *Arabidopsis* flower.

## Results

### Single cell transcriptomics confirms increased boundary cell identity and suggests decreased translation is cell type specific in young *drmy1* flower buds

To enrich for flower buds early in sepal development, we introgressed the *drmy1-2* mutation into the *apetala1 cauliflower* 35S::APETALA1-GR (*ap1cal* 35S::AP1-GR) background, which produces large cauliflower-like clusters of densely packed arrested floral meristems (Figure S1A-S1B) (Wellmer et al. 2006; Kong et al. 2024b). We induced floral initiation with dexamethasone and harvested tissue 108 hours later when flower buds spanned from developmental stage 2 (no sepal primordia) to late stage 3 (all sepals initiated and beginning to elongate) (Smyth et al. 1990) (Figure S1A). We prepared protoplasts from two independent biological replicates per genotype [*ap1cal* 35S::AP1-GR (“WT”) and *drmy1 ap1cal* 35S::AP1-GR (“*drmy1*”)] using a meristem optimized protocol (Satterlee et al. 2020) and performed 10x Genomics scRNA-seq. After filtering, we obtained 93,365 high quality cells (mean of 6457 unique molecular identifiers and 2613 features (genes) per cell); (Figure S1C) and found minimal inter-replicate variability (Figure S1D), allowing us to combine replicates by genotype for downstream analysis.

Principal component analysis (PCA) initially clustered many cells by cell-cycle phase, even after cell cycle regression was performed (Figure S1E), reflecting active cell division in floral meristems. To fully remove the effects of cell cycle on downstream analysis, we excluded genes where >3% of variance was explained by cell cycle phase (Supplementary Data 4). This filtering yielded 17 transcriptionally distinct clusters (Figure 1B), which we then annotated using established marker genes (Figure 1C). Broadly, clusters 1, 10, 12, and 13 represent epidermal cells and express epidermal markers such as ATML1 and PDF1 (Figure 1C). Clusters 0, 1, 2, 4, 5, 8, and 9 are largely meristematic cells, expressing markers such as STM, AG, WUS, CLV, and others (Figure 1C). Clusters 3 and 15 represent mesophyll cells and have a relatively large proportion of transcripts from chloroplast genes (Figure 1C, Figure S1I). Clusters 7, 11, and 15 express vasculature specific genes such as APL and TDR (Figure 1C). Of particular interest to us for understanding the development of young flower buds are clusters 1 (meristematic epidermis), 10 (mature epidermis), 0 (core meristem), and 8 (organ primordia).

Additionally, we aimed to identify boundary cells, given that prior work established overexpression of boundary identity as a key factor in *drmy1* floral development (Kong et al. 2024c). Particularly, CUC1 protein and transcriptional reporters have higher expression levels and expanded expression domains in *drmy1* (Kong et al. 2024c). However, canonical boundary genes such as the *CUC* genes were sparsely expressed in our scRNA-seq dataset. To identify these cells, we used a pipeline developed by Neumann et al. (2022) that maps scRNAseq clusters onto a 3-dimensional floral meristem atlas based on shared gene expression patterns. Accuracy of this approach was confirmed by several single cell clusters which mapped with high probability to their expected locations on the floral meristem atlas. This included cluster 0 (core meristem) to the core of the atlas; cluster 1 (meristematic epidermis) to the L1 layer of the atlas; and cluster 8 (primordia) to the sepal primordia (Figure 1D). Cluster 2 mapped to the boundary between the core meristem and the initiating primordia, indicating probable boundary cells for further analysis (Figure 1D).

Interestingly, cluster 2 in *drmy1* mapped to the boundary region with higher probability than in WT (0.22 in *drmy1* vs 0.14 in WT) (Figure 1D). We simultaneously see that cluster 2 (boundaries) represents a much higher proportion of the total cell population in *drmy1* (0.125) than in WT (0.087) (Figure 1E), which may reflect the upregulation of boundary identity in *drmy1*. Other clusters with large differences in cell population proportions include fewer cells in cluster 3 (mesophyll) and more cells in cluster 4 (carpels) in *drmy1* (Figure S1F). This is likely because *drmy1 ap1cal* 35S::AP1-GR cauliflower-like clusters have fewer leafy bracts and more flower buds that prematurely develop before dexamethasone induction (Figure S1B).

To study differential gene expression patterns in *drmy1*, we performed differential gene expression analysis and gene ontology (GO) term enrichment analysis on pseudobulked scRNA-seq data. Terms related to translation, ribosomes, and RNA binding were greatly enriched in WT compared to *drmy1* (Figure 1F) as seen previously in Kong et al. (2024b). Notably however, this enrichment was much lower or absent in clusters 0 (core meristem), 1 (meristematic epidermis), 10 (mature epidermis), 2 (boundaries), 12 and 13 (guard cells) (Figure 1F). This suggests that decreases in translation may not be as global in *drmy1* than previously thought, and perhaps some cell types are more robust to changes in translation than others. In conclusion, our scRNA-seq dataset allowed us to identify key cell types present in developing flower buds while also recapitulating known trends in boundary cell identity upregulation and translation defects in *drmy1*. Our next step was to test if this dataset would allow us to see transcriptomic changes in auxin and cytokinin signaling not apparent from bulk RNAseq, as well as search for novel transcriptomic changes in *drmy1*.

## ScRNA-seq recapitulates patterns of decreased auxin signaling, increased cytokinin signaling, and reveals increased brassinosteroid signaling in *drmy1*

To look for transcriptomic changes in hormone signaling pathways, for each cell we calculated the percentage of transcripts from genes up or down regulated by the application of various hormones based on a study by Nemhauser et al. (2006) out of the total number of transcripts in the same cell. We expect that the signaling levels of a given hormone will be positively correlated with the percentage of transcripts from genes upregulated by that hormone, and negatively correlated with genes downregulated by that hormone (Figure 2A). This can be used to infer signaling levels of various hormones in each cell. For instance, if auxin signaling is higher in WT than *drmy1* as seen in Kong et al. (2024b), we expect the transcripts from auxin upregulated genes to be more abundant, and those of downregulated genes to be less abundant, in WT relative to *drmy1* (Figure 2A). This is exactly the trend we observe in our data, suggesting that auxin signaling is higher in WT, especially in clusters 0 (core meristem), 10 (mature epidermis), and 3 (mesophyll) (Figure 2B). We see the opposite trend with cytokinin regulated genes – particularly with lower expression of cytokinin downregulated genes in *drmy1*, suggesting cytokinin signaling is higher in *drmy1* (Figure 2B). This trend is especially apparent in clusters 0 (core meristem), 1 (meristematic epidermis), 10 (mature epidermis), and 3 (mesophyll) (Figure 2B).

**Figure 2:**
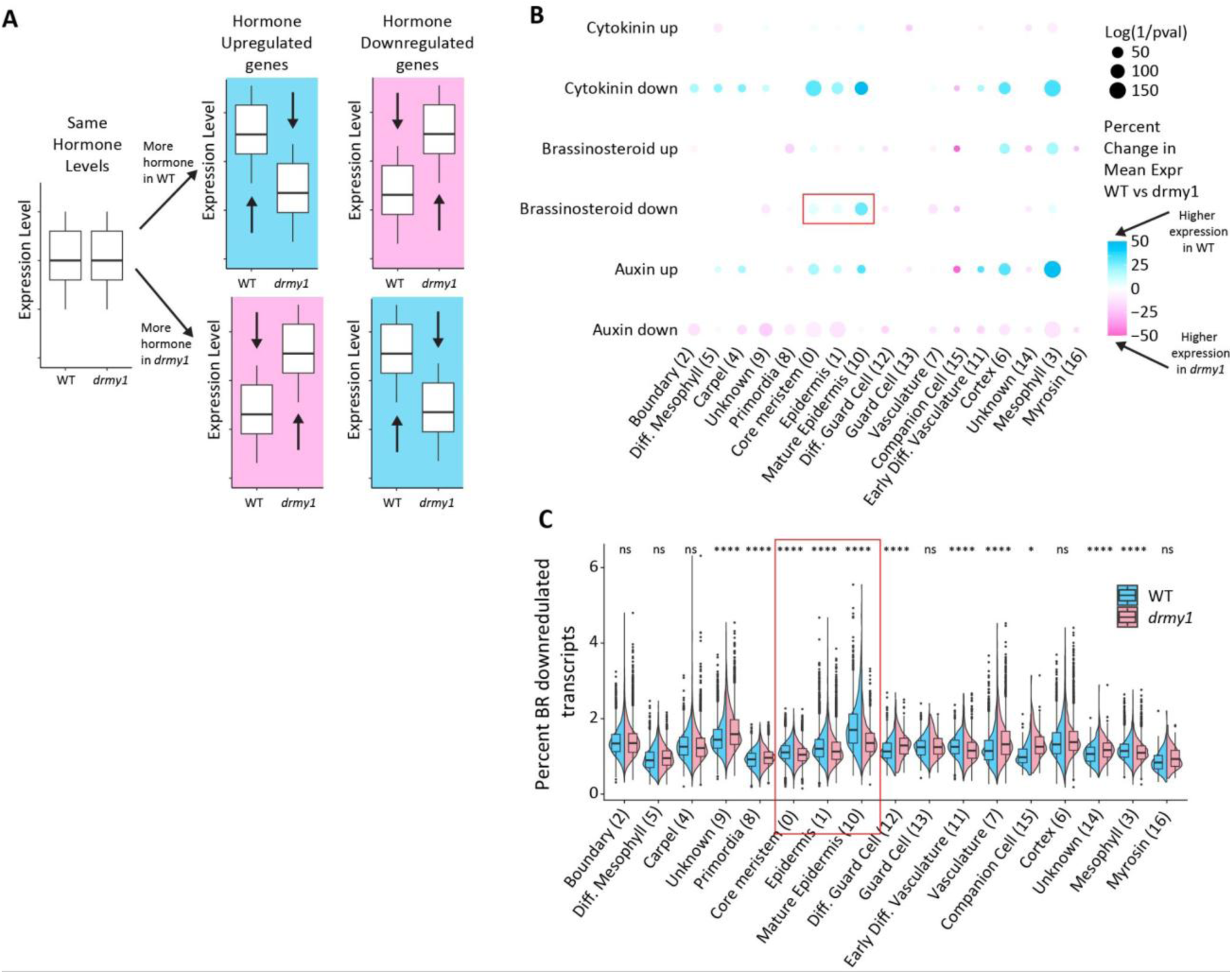
Gene expression trends are consistent with decreased cytokinin signaling, and increased auxin and brassinosteroid signaling in *drmy1*. A) Schematic of the hypothesized changes in expression levels of hormone up and down-regulated genes under high and low levels of hormone in WT or *drmy1*. B) Dotplot showing the differences in expression of genes up and down-regulated by auxin, cytokinin, and brassinosteroids. For each cell, the percentage of transcripts that come from the set of genes up or downregulated by applications of these hormones according to Nemhauser et al. (2006) is calculated, then the mean of all the cells in each cluster is calculated for WT and *drmy1*. The size of each dot represents the log of the inverse of the p-value between the mean of the two genotypes with a Bonferroni adjusted threshold of p < 0.000245. The color of each dot represents the percentage increase or decrease of the mean of *drmy1* compared to WT. Red box indicates changes in BR signaling in clusters of interest in young flower buds (core meristem, epidermis, and mature epidermis). See figure S2 for other hormones. C) Violin plot showing more details of the change in expression patterns from the “Brassinosteroid down” row from panel B. Red box corresponds to the same data boxed in panel B. *p < 0.00294, **p < 0.00059, *** p < 0.000059, **** p < 0.0000059 Bonferroni adjusted p-value thresholds in Wilcoxon’s rank sum tests compared with WT. See also: Figure S2, Supplementary Data 2

After confirming known changes to auxin and cytokinin signaling, we performed the same analyses on other hormones as well, especially looking for changes in the core meristem, epidermal, and boundary clusters. Genes involved in brassinosteroid (BR) signaling stood out, especially in the core meristem and epidermal clusters where BR downregulated genes have lower expression in *drmy1* (Figure 2B and 2C), which suggests possible BR upregulation in *drmy1*. We also looked at any individual gene differentially expressed that had the word “brassinosteroid” in its GO terms. Notably, BR biosynthesis gene *DWF1* is upregulated in *drmy1* while *DWF4*, which has a negative feedback loop with BR signaling (Kim et al. 2006), is downregulated (Figure S2B). BSK2, a core component of the BR signaling pathway, is also upregulated in *drmy1* (Figure S2B). We performed similar analyses with other hormones such as abscisic acid, jasmonic acid, and gibberellic acid, and while there were notable changes in their expression patterns in several clusters, the overall trends were less clear in our clusters of interest, thus we focused primarily on BR going forward (Figure S2A). These results suggest that BR signaling may be higher in *drmy1*, thus we decided to investigate brassinosteroids further.

### Visium spatial RNAseq corroborates scRNAseq trends with brassinosteroids and boundaries

While single cell RNAseq offers excellent resolution, it does not preserve spatial information. Therefore, to understand the spatial patterns of gene expression changes in *drmy1*, we applied Visium Spatial RNAseq to 10 μm sections of floral meristem clusters from WT and *drmy1* in *ap1cal* 35S::AP1-GR backgrounds 108 hours post induction, when numerous flower buds on the cauliflower-like head are starting to initiate sepals. These transverse slices allow for easy visual identification of many common cell types and anatomical structures (Figure 3A).

**Figure 3:**
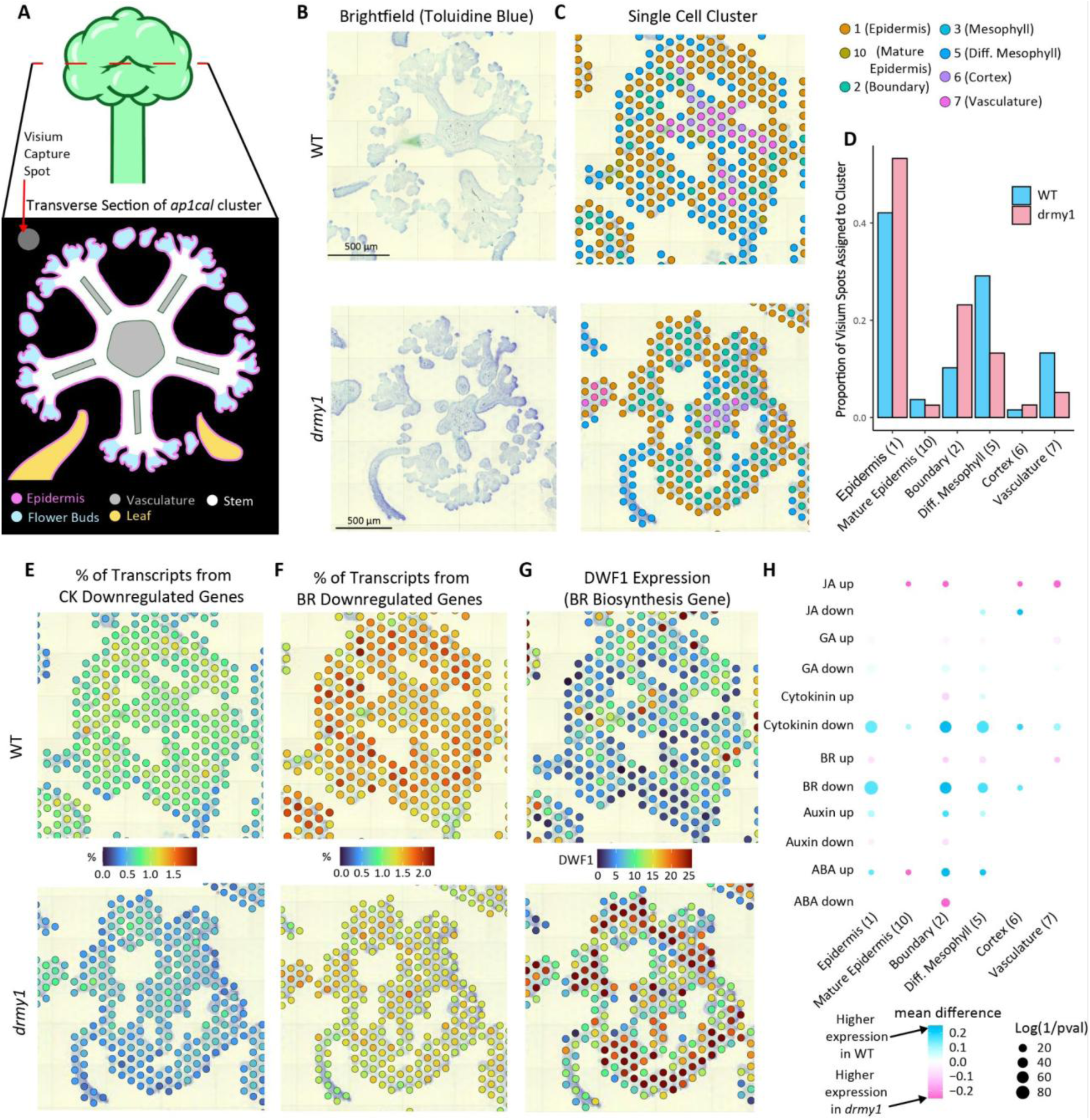
Spatial RNAseq further suggests boundary identity, cytokinin signaling, and brassinosteroid signaling are increased in *drmy1*. A) Cartoon of a typical transverse section of an *ap1cal* 35S::AP1-GR cauliflower-like cluster, showing the developing flower buds with initiating sepals on the outer ring . B) Representative brightfield images of transverse sections of WT and *drmy1* in the *ap1cal* 35S::AP1-GR background. n=9 Total WT replicate cauliflower-like clusters, n=22 *drmy1* replicate cauliflower-like clusters (See also Figure S3). C) An overlay of which single cell cluster maps with the highest prediction score to each Visium spot on the same tissue shown in panel (B). See figure S3 for a more detailed view of prediction scores from each single cell cluster. D) The proportions of Visium spots grouped by which single cell cluster mapped to each with the highest prediction score (clusters that mapped to <10 spots in either genotype were omitted). For cluster 2 (boundaries), WT = 0.102, *drmy1* = 0.232. E-G) Overlays of the percentage of transcripts from cytokinin downregulated genes (E), brassinosteroid downregulated genes (F), and SCT normalized expression of *DWF1*, a brassinosteroid biosynthesis gene (G). H) Dotplot showing the differences in expression of genes up and down-regulated by various hormones. For each cell, the percentage of transcripts that come from the set of genes up or downregulated by applications of these hormones according to Nemhauser et al. (2006) is calculated, then the mean of all the cells in each cluster is calculated for WT and *drmy1*. The size of each dot represents the log of the inverse of the p-value between the mean of the two genotypes with a Bonferroni adjusted threshold of p < 0.00069. The color of each dot represents the percentage increase or decrease of the mean of *drmy1* compared to WT. Scale bars 500 μm See also: Figure S3, S4

Although the Visium capture spots were slightly too large (55 μm diameter spots placed 100 μm apart) to compare gene expression profiles from different regions of individual flower buds (about 100 μm in diameter), we were still able to use Seurat Deconvolution to predict the cell type composition of the spots based on our single cell clusters (Figure 3B-3D, S3B). In agreement with our scRNAseq results, boundary cell identity (cluster 2) had the highest prediction score in a higher proportion of Visium spots in *drmy1* (0.232) than in WT (0.102) (Figure 3C and 3D). Fitting with the morphology, the boundary identity (cluster 2) maps best to spots just interior to epidermis-mapping (cluster 1) spots on the edges of floral meristem clusters (Figure 3C).

To clarify the spatial distribution of hormone signaling, we summarized the difference in the composition of sets of genes up and downregulated by various hormones between WT and *drmy1*. We find that cytokinin and BR downregulated genes have much higher expression in WT, especially in spots whose prediction scores are highest for the epidermis, boundary, and mesophyll identities (Figure 3H). This further supports the hypothesis that cytokinins and BR are upregulated in *drmy1*, especially in spots on and nearby to young flower buds (Figure 3E-3G).

Similarly, key BR genes such as the BR biosynthesis gene *DWF1* are highly upregulated in *drmy1* spots on and near the flower buds (Figure 3H). Together, these results verify that brassinosteroid signaling may be upregulated in *drmy1* and is a good candidate for further validation and exploration of sepal elongation defects.

### A ratiometric BES1 reporter confirms brassinosteroid signaling is upregulated in *drmy1*

To confirm that BR signaling is upregulated in *drmy1*, we introgressed the ratiometric BRI1-EMS-SUPPRESSOR1 (BES1) reporter (*pUBQ10::BES1-ypet/pUBQ10::H2B-mCherry*) developed by Ackerman-Lavert et al. (2021) into *drmy1*. Compared to WT, *drmy1* sepals had significantly higher mean BES1/H2B ratios (p = 0.0041) (Figure 4A and 4B). When broken down by sepal position, we find that BR signaling was significantly higher in *drmy1* inner and lateral sepals, while in outer sepals the difference is insignificant but trends higher (outer; p = 0.15, inner; p = 0.025, lateral; p = 0.048) (Figure 4C). In addition, the coefficient of variance (CV) of BES1/H2B ratios within individual sepals is significantly higher in *drmy1* (p = 6.5 x 10^-12^) (Figure 4D), which is consistent with *drmy1* having increased variability in auxin signaling as well (Kong et al. 2024c).

**Figure 4:**
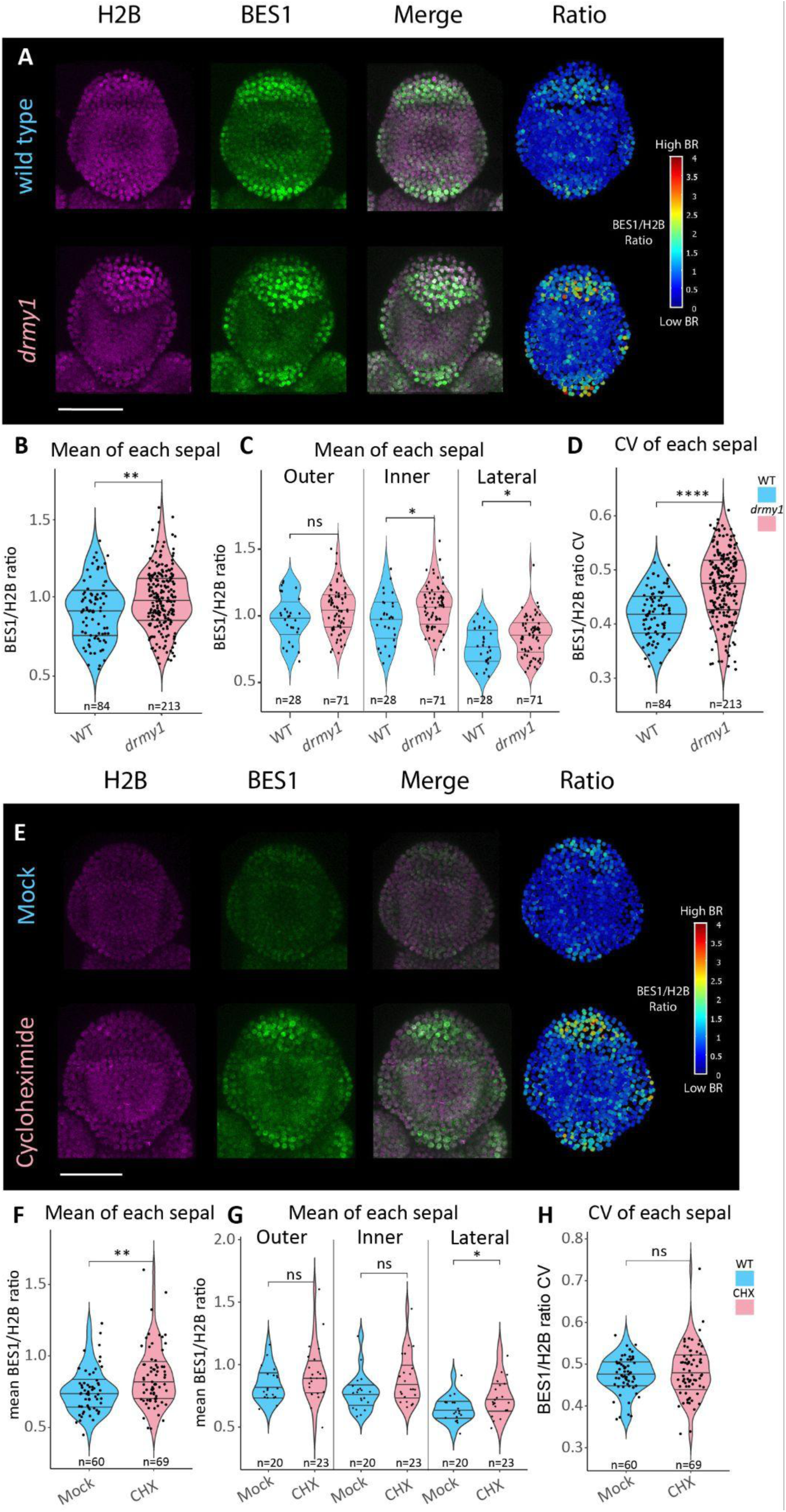
A BES1 ratiometric reporter confirms brassinosteroid signaling is upregulated in *drmy1*. A) WT and *drmy1* expressing *pUBQ10::H2B-mcherry* (magenta)/*pUBQ10::BES1-ypet* (green), both channels merged, and the ratio of the BES1 signal divided by the H2B signal in each nucleus. B) Quantification of the mean BES1/H2B signal ratio for each sepal, p = 0.0041. C) Quantification from (B) separated by sepal position, p = 0.15 (outer), p = 0.025 (inner), p = 0.048 (lateral). D) CV of BES1/H2B ratio in each sepal, p = 6.5 x 10^-12^. E) Day 6 mock and 2 μM cycloheximide treated buds expressing the BES1/H2B ratiometric reporter. F) Quantification of the mean BES1/H2B signal ratio for each sepal, p = 0.0049. G) Quantification from (F) separated by sepal position, p = 0.18 (outer), p = 0.059 (inner), p = 0.03 (lateral). H) CV of BES/H2B ratio in each sepal, p = 0.62. Lines in violin plots represent the 25^th^, 50^th^, and 75^th^ quantiles. For each graph, n is listed below the violin plot. *p < 0.05, **p < 0.005, *** p < 0.0005, **** p < 0.00005 in Wilcoxon’s rank sum tests. Scale bars 50 μm

Previous work has shown that treating WT buds with the translation inhibitor cycloheximide (CHX) can recreate several phenotypes seen in *drmy1,* such as sepal growth defects and changes to auxin and cytokinin signaling (Kong et al. 2024b). Therefore we hypothesized that we should be able to recreate increases in brassinosteroid signaling with CHX as well. As anticipated, treating WT plants with 2 μM CHX for 6 days significantly increased BES1/H2B ratios of sepals (p = 0.0049) (Figure 4E and 4F). This result was only significant for lateral sepals, though trended higher in outer and inner sepals (outer; p = 0.18, inner; p = 0.059, lateral; p = 0.03) (Figure 4G). However, CHX did not recapitulate the increase in CV seen in *drmy1* (Figure 4H), suggesting that the increased variability in BR signaling in *drmy1* is not a direct consequence of the decreased translation. Together, these results confirm that brassinosteroid signaling is increased in *drmy1* likely due to the decrease in protein translation.

### Altering brassinosteroid levels differentially affects elongation of the inner and outer sepals

Because *drmy1* has increased BR signaling, we investigated BR mutants for sepal growth defects. We noticed that the dominant *brassinazole-resistant1-1D* (*bzr1-1D*) mutant, which has higher levels of BR signaling due to a stabilized BR responsive transcription factor, has elongated outer sepals compared to WT (p = 0.019), reminiscent of the elongated outer sepals of *drmy1* (Figure 5A and 5B). Interestingly, *bzr1-1D* inner sepals are not longer than WT inner sepals (p = 0.87) (Figure 5C). This results in a larger discrepancy between the lengths of the inner and outer sepals in *bzr1-1D* compared to WT, as quantified by taking the ratio of the lengths of the outer divided by the inner sepals (p = 0.00014) (Figure 5D). However, the outer/inner ratio is not quite as severe in *bzr1-1D* as in *drmy1*, suggesting that increased BR signaling accounts for some, but not all, of the size discrepancy between the outer and inner sepals in *drmy1* (Figure 5D).

**Figure 5:**
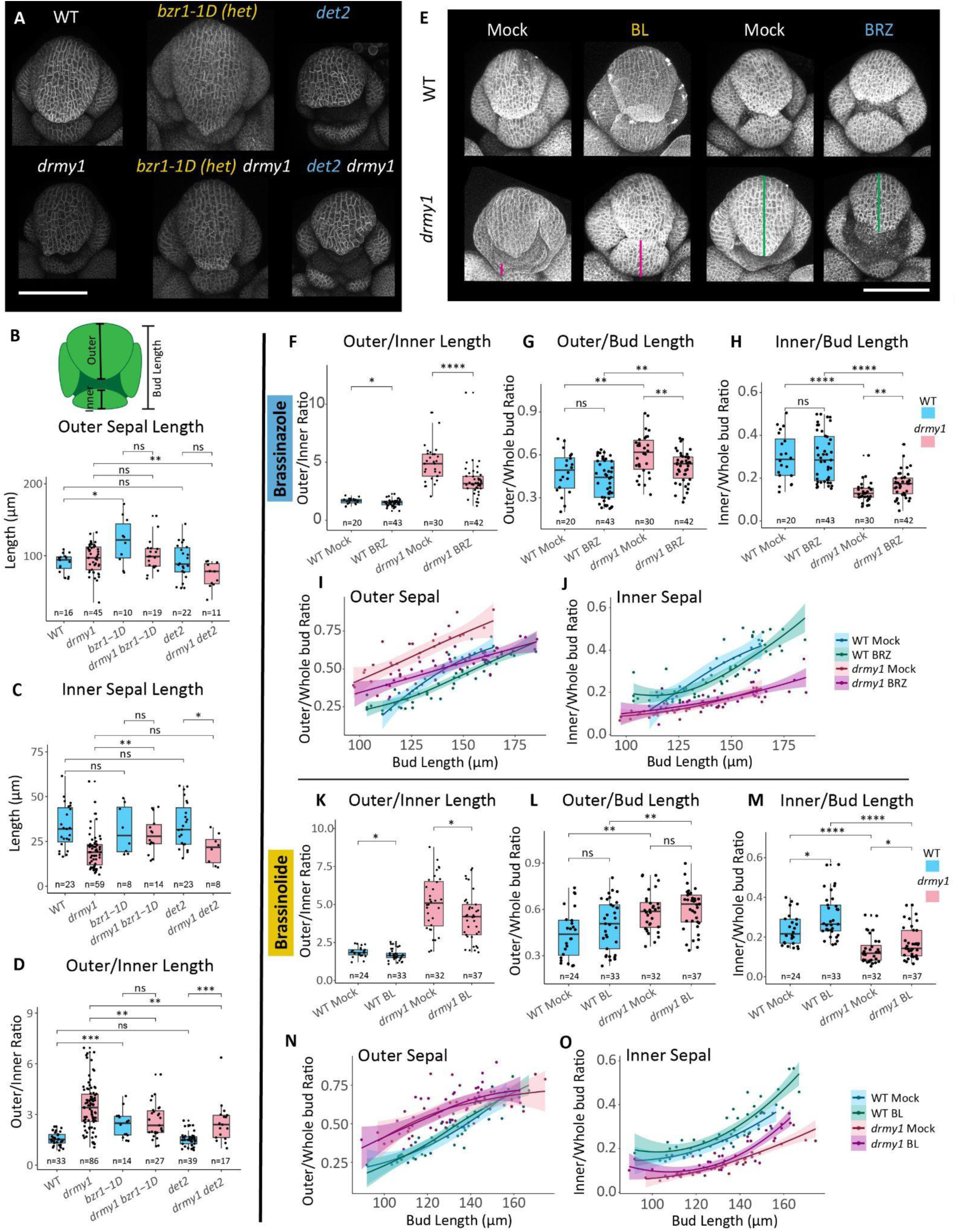
Sepal growth defects are partially rescued in *drmy1* by decreasing brassinosteroid signaling to reduce outer sepal overgrowth, or by increasing brassinosteroid signaling to increase inner sepal growth. A) Confocal images of single brassinosteroid mutants *bzr1-1D* and *det2*, as well as double mutants with *drmy1* expressing the membrane marker *35S::RCI2A-mCitrine*. B – D) Quantification of the lengths of the outer sepals of stage 5-7 flower buds (B), inner sepals of stage 3 – 5 flower buds (C), and the ratio between the length of the outer sepal divided by the length of the inner sepal in all flower buds (D) in single and double mutants. (B) WT vs *bzr1-1D* p = 0.019; WT vs *det2* p = 0.87; *drmy1* vs *drmy1 bzr1-1D* p = 0.93; *drmy1* vs *drmy1 det2* p = 0.0018; *bzr1-1D* vs *drmy1 bzr1-1D* p = 0.13; *det2* vs *drmy1 det2* p = 0.069. (C) WT vs *bzr1-1D* p = 0.87; WT vs *det2* p = 0.73; *drmy1* vs *drmy1 bzr1-1D* p = 0.0017; *drmy1* vs *drmy1 det2* p = 0.44; *bzr1-1D* vs *drmy1 bzr1-1D* p = 0.71; *det2* vs *drmy1 det2* p = 0.013. (D) WT vs *bzr1-1D* p = 0.00014; WT vs *det2* p = 0.84; *drmy1* vs *drmy1 bzr1-1D* p = 0.0047; *drmy1* vs *drmy1 det2* p = 0.0045; *bzr1-1D* vs *drmy1 bzr1-1D* p = 0.45; *det2* vs *drmy1 det2* p = 0.00025. E) Confocal images of WT and *drmy1* flower buds treated for 6 days with either 50 uM brassinazole (BRZ), a brassinosteroid biosynthesis inhibitor, 400 nM brassinolide (BL), or mock treatments. Buds are either expressing the membrane marker *35S::RCI2A-mCitrine* or are stained with 0.1 mg/mL propidium iodide. Note magenta bars indicating rescued growth of the inner sepal in BL treated *drmy1* buds, and the red bars indicating reduced overgrowth of outer sepals in BRZ treated *drmy1* buds. F-H) Quantification of the ratio between the lengths of the outer and inner sepals (F), the ratio between the lengths of the outer sepal and the entire bud (G), and the ratio between the lengths of the inner sepal and the entire bud (H) after mock or BRZ treatment. The length of the entire bud is an indication of the developmental stage since the bud increases size as it grows. (F) WT mock vs BRZ p = 0.032; *drmy1* mock vs BRZ p = 1 x 10^-5^. (G) WT mock vs BRZ p = 0.25; *drmy1* mock vs BRZ p = 0.0094; WT mock vs *drmy1* mock p = 0.0057; WT BRZ vs *drmy1* BRZ p = 0.0094. (H) WT mock vs BRZ p = 0.95; *drmy1* mock vs BRZ p = 0.0046; WT mock vs *drmy1* mock p = 3.4 x 10^-9^; WT BRZ vs *drmy1* BRZ p = 5.6 x 10^-8^. I-J) Relationship between the length of the entire bud and the ratio between the lengths of the outer sepal (I) or the inner sepal (J) after mock or BRZ treatment. Lines represent the Loess regression fit of the data and shaded areas represent the 95% confidence interval. K-M) Quantification of the ratio between the lengths of the outer and inner sepals (K), the ratio between the lengths of the outer sepal and the entire bud (L), and the ratio between the lengths of the inner sepal and the entire bud (M) after mock or BL treatment. (K) WT mock vs BL p = 0.025; *drmy1* mock vs BL p = 0.035. (L) WT mock vs BL p = 0.16; *drmy1* mock vs BL p = 0.33; WT mock vs *drmy1* mock p = 0.0014; WT BL vs *drmy1* BL p = 0.0094. (M) WT mock vs BL p = 0.026; *drmy1* mock vs BL p = 0.044; WT mock vs *drmy1* mock p = 1.5 x 10^-6^; WT BL vs *drmy1* BRZ p = 6.1 x 10^-7^. N-O) Relationship between the length of the entire bud and the ratio between the lengths of the outer sepal (I) or the inner sepal (J) after mock or BL treatment. Lines represent the Loess regression fit of the data and shaded areas represent the 95% confidence interval. *p < 0.05, **p < 0.005, *** p < 0.0005, **** p < 0.00005 in Wilcoxon’s rank sum tests. Scale bars 100 μm See also: Figure S5

We next wondered if reduced BR levels would alter the relative elongation of the outer and inner sepals flower buds. For this we used the *de-etiolated2* (*det2*) mutant, which has greatly reduced levels of brassinolide biosynthesis. Interestingly, *det2* does not significantly alter the lengths of the inner (p = 0.73) or outer (p = 0.87) sepals compared to WT, and thus does not affect the outer/inner sepal ratio (p = 0.84) (Figure 5A-5D).

Next we wanted to know if crossing *drmy1* with these BR mutants could rescue or worsen the sepal elongation defects. We hypothesized that further increasing BR signaling by crossing *drmy1* with *bzr1-1D* would accentuate outer sepal overgrowth, but were surprised to find that outer sepal length remained unchanged in *drmy1 bzr1-1D* double mutant compared to *drmy1* alone (p = 0.93), suggesting the outer sepal growth induced by BR signaling is already saturated (Figure 5A and 5B). Instead, we noticed that the inner sepal length actually increased in the double mutant (p = 0.0017) (Figure 5A and 5C), leading to a partial rescue of the outer/inner sepal ratio (p = 0.0047) (Figure 5D), suggesting that further increasing BR signaling can promote elongation of the inner sepal.

Since BR signaling is upregulated in *drmy1*, we hypothesized that crossing *det2* with *drmy1* may rescue the sepal primordium size coordination defect in *drmy1* by reducing outer sepal overgrowth. Compared with *drmy1*, the *drmy1 det2* double mutant has reduced outer sepal length (p = 0.0018) while inner sepal length remains unchanged (p = 0.44) (Figure 5A-5C). As a result, the outer/inner sepal ratio is partially rescued in *drmy1 det2* compared to *drmy1* (p = 0.0045) (Figure 5D). These results suggest that decreasing BR signaling levels can partially rescue the *drmy1* sepal primordium size coordination defect by limiting the overgrowth of the outer sepal.

To verify the trends from our mutant analysis, we altered BR signaling through chemical treatments as well. We achieved this by applying 400 nM brassinolide (BL), the most bioactive brassinosteroid, or 50 μM brassinazole (BRZ), an inhibitor of DWF4, a key enzyme in the BR biosynthesis pathway. Dipping inflorescences in solutions of these chemicals daily for 6 days clearly altered tissue growth patterns in expected ways, such as increasing (in the case of BL) or decreasing (in the case of BRZ) pedicel and silique lengths (Figure S5A). As predicted from our mutant experiments, both BL and BRZ applications lead to a partial rescue of sepal primordium size coordination as represented by the outer/inner sepal ratios in *drmy1* (BRZ p = 1 x 10^-5^; BL p = 0.035) (Figure 5E, 5F and 5K). Interestingly, both chemicals also lead to small but significant decreases in the outer/inner sepal ratios in WT (BRZ p = 0.032; BL p = 0.025) (Figure 5E, 5F and 5K), but neither inner nor outer sepals individually were significantly affected by BRZ (outer p = 0.25; inner p = 0.95) (Figure 5G and 5H). As anticipated, BRZ significantly decreases the length of the outer sepal normalized to the length of the whole bud in *drmy1* (p = 0.0094) (Figure 5G and 5I). BRZ has no obvious effect on inner sepal relative length in *drmy1* (Figure 5J). Altogether, these results corroborate the idea that decreasing BR levels can reduce outer sepal overgrowth in *drmy1* without affecting inner sepal growth.

Conversely, BL did not lead to any significant changes in outer sepal size normalized to the whole bud in *drmy1* (p = 0.33) (Figure 5L and 5N). Instead, BL treatments significantly increased the size of the inner sepal normalized to the whole bud in both WT and *drmy1* (WT p = 0.026; *drmy1* p = 0.044) (Figure 5M). This effect was more pronounced at later stages of growth, when buds are larger, possibly because later stages have had more time to accumulate more growth (Figure 5O).

From prior work, we know that a large contributor to the defects in *drmy1* sepal development is the variable and delayed sepal initiation caused by decreased auxin signaling and increased CUC1 expression (Zhu et al. 2020, Kong et al. 2024c). To ensure the partial rescue from BRZ and BL treatments is a result of changes to BR mediated elongation and not simply due to rescuing sepal initiation, we imaged treated plants every 6 hours and recorded when sepals initiated. In doing so, we found that the treatments did not rescue sepal initiation timing in *drmy1* (Figure S5B). Furthermore, to rule out cross talk between BR and auxin or CUC1, we applied these treatments to *DR5::Venus-N7* and to plants expressing *pCUC1::3xVenus-N7*, and found no obvious changes to the expression intensity or pattern of either reporter (Figure S5C and S5D).

These results confirm that increasing BR levels partially rescues *drmy1* sepal primordium elongation defects by increasing the growth of the inner sepal, while decreasing BR levels partially rescues by decreasing the overgrowth of the outer sepal, and that these effects are independent of the defects in sepal initiation caused by auxin and CUC1.

### Altering brassinosteroid levels alters relative sepal sizes by changing outer and inner sepal growth rates

To better understand the impact of altering BR levels on growth patterns of the different sepals, we live imaged WT and *drmy1* with BRZ and BL treatments. We normalized for differences in overall growth rates between buds within and between treatments by calculating the total change in area of all sepals, and then determining the proportion of the total change in area that comes from each sepal. In mock treated WT buds, the outer sepal accounts for a slight majority of growth between each timepoint (typically 50-60%), while the inner sepal accounts for less (20-30%) of the growth, leading to relatively small outer/inner sepal ratios (Figure 6C and 6D). Treatment with BL increases the relative growth of the outer sepal, particularly at earlier timepoints (60-70%), leading to slightly larger outer/inner sepal ratios (Figure 6A and 6B). Meanwhile, BRZ slightly decreases the growth contributions of the outer sepal, particularly at later timepoints (40-50%), although the effect is small enough that the outer/inner sepal is largely unchanged (Figure 6E and 6F). Additional replicates show similar trends (Figure S6).

**Figure 6:**
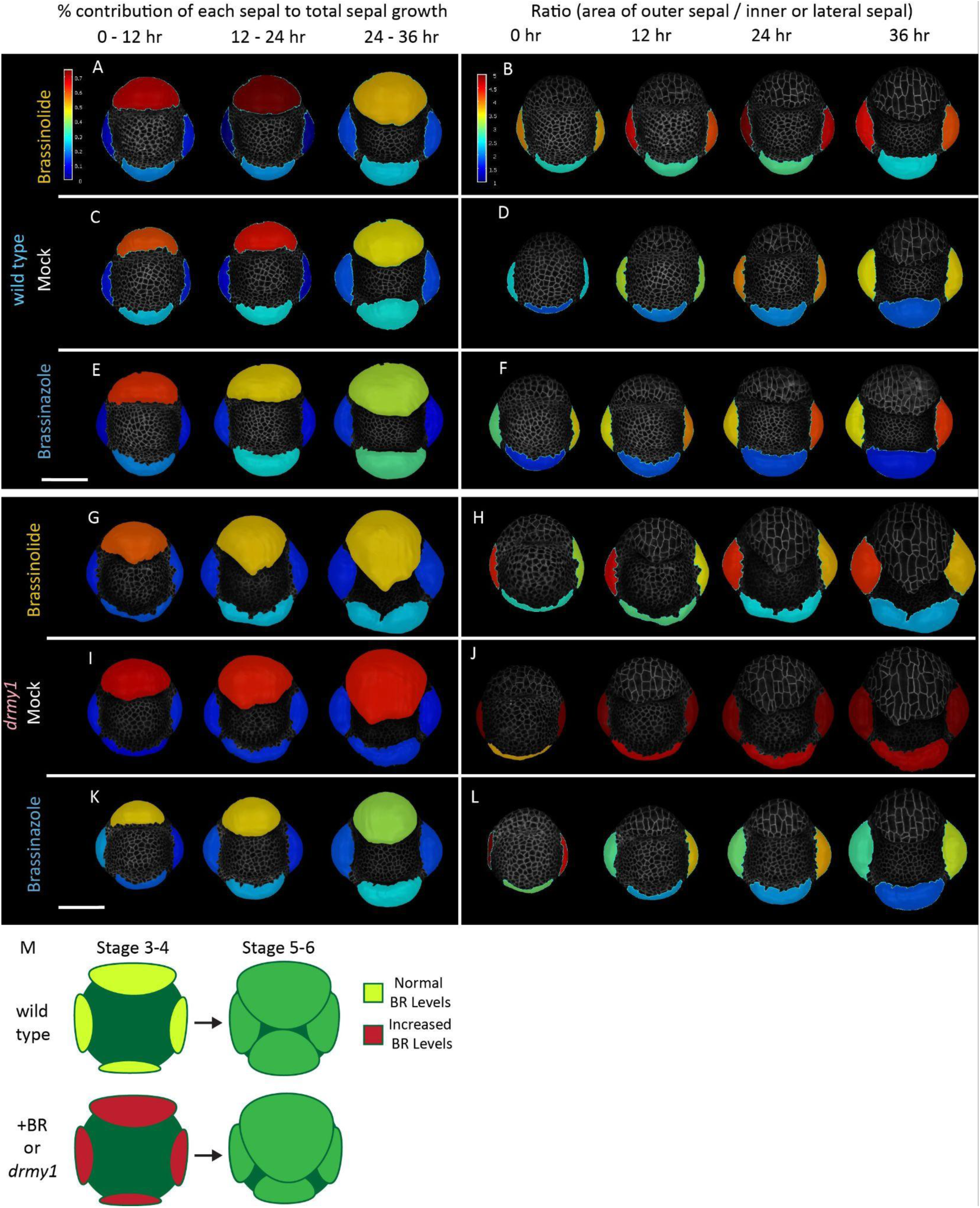
Brassinazole reduces outer/inner sepal ratios by reducing outer sepal growth, brassinolide by increasing inner sepal growth. A, C and E) Proportion of total sepal growth contributed by each sepal in WT buds treated with 400 nM brassinolide (BL) (A), mock (C), or 50 μM brassinazole (BRZ) (E) at 12 hour time intervals. B, D, and F) Corresponding ratio between the area of the outer sepal to the area of the inner and each lateral sepal under BL (B), mock (D), and BRZ (F) treatments in WT buds. G, I and K) Proportion of total sepal growth contributed by each sepal in *drmy1* buds treated with 400 nM brassinolide (BL) (A), mock (C), or 50 uM brassinazole (BRZ) (E) at 12 hour time intervals. H, K and L) Corresponding ratio between the area of the outer sepal to the area of the inner and each lateral sepal under BL (B), mock (D), and BRZ (F) treatments in *drmy1* buds. M) Model representing the role of increased BR signaling or the *drmy1* mutation on the growth of the *Arabidopsis* flower buds. Scale bars 50 μm See also: Figure S6, S7, S8

Also, segmenting every cell in the sepal shows that these differences in growth rates are driven largely by cells in the sepal tips, which are the fastest growing cells at these early stages (Figure S8G – S8I). These results recapitulate trends seen in our single timepoint images, which demonstrate that BL can increase outer sepal growth and BRZ can decrease outer sepal growth in WT, but that growth remains mostly robust.

In contrast, outer sepals from mock treated *drmy1* buds account for a much larger proportion of sepal growth (>70%), leading to very high outer/inner sepal ratios (Figure 6I and 6J). Applying BL reduces outer sepal growth percentage to near WT levels (50-60%) and increases inner sepal growth percentage (20-30%), greatly reducing the outer/inner sepal ratios (Figure 6G and 6H). Furthermore, absolute growth amounts support that the amount of growth in the outer sepal is similar between BL and mock, but inner sepal growth has increased (Figure S8D and S8E). BRZ treated buds have greatly reduced outer sepal growth contributions (40-50%) (Figure 6K and 6L), while absolute growth rates confirm that the inner sepal remains relatively unaffected (Figure S8F). Additional replicates show similar trends, and intriguingly, the severity of the growth defect as visualized by outer/inner sepal ratio is worse in samples where outer sepal growth contributions are higher (Figure S7). Our results confirm that increasing BR levels in *drmy1* partially rescues sepal elongation defects by increasing inner sepal growth and not altering outer sepal growth, while decreasing BR levels partially rescues by decreasing outer sepal growth rates without altering inner sepal growth rates (Figure 6M).

## Discussion

Growth coordination is an essential facet of developmental robustness. By profiling transcriptomic changes at the single cell level in *drmy1*, we are able to show that brassinosteroids, which are key regulators of cellular growth and elongation, are upregulated in *drmy1* during early sepal elongation. An important defect hindering robust sepal development in *drmy1* is the lack of proper coordination between the elongation of the outer and inner sepals.

We use BR mutants to demonstrate the upregulation of BR signaling can lead to overgrowth of the outer sepal in WT flowers (Figure 5B). This is similar to the overgrowth defect seen in *drmy1*, which demonstrates that proper BR signaling levels are required for coordination of outer and inner sepal growths (Figure 6M). We also use brassinolide and brassinazole treatments to show that increasing BR signaling in *drmy1* can partially rescue the sepal size discrepancies by increasing growth of the inner sepal, while decreasing BR signaling partially rescues by decreasing overgrowth of the outer sepal (Figure S5E).

The differential effects of BR signaling on outer and inner sepal primordium elongation seen here are not entirely surprising given that previous research has shown BR can have different effects on different populations of cells or tissues (Singh and Savaldi-Goldstein 2015). For example, in rice leaves BR preferentially inhibits cell proliferation on the abaxial side of the lamina joint, causing changes in leaf erectness (Sun et al. 2015). BR may also have differential effects on the epidermis compared to interior cell layers and may act predominantly through the epidermis (Hacham et al. 2011; Savaldi-Goldstein et al. 2007; Zhiponova et al. 2012), although this finding is controversial and may reflect low levels of receptor expression in other cell layers (Blanco-Touriñán et al., 2024).

Previous studies of BR in the shoot apical meristem have shown that downregulation of BR in boundary cells is important for proper *CUC* gene expression and boundary formation (Tsuda et al. 2014; Gendron et al. 2012). It is surprising, then, that despite a clear upregulation of boundary gene identity in *drmy1* (Figure 1E; Figure 3D) (Kong et al. 2024c), that BR signaling is still overall higher in *drmy1* (Figure 3A-B). However a closer look at the boundary cell type in our single cell data shows that BRs are in fact not upregulated in the boundaries (cluster 2) (Figure 2A).

Intriguingly, the BR treatments in *drmy1* do not appear to alter auxin signaling mediated sepal initiation, suggesting that the partial rescue of sepal elongation is not a direct consequence of BR – auxin crosstalk. However, the interplay between these hormones is often very complex and changes depending on the circumstances and location (Halliday 2004; Nemhauser et al. 2004). In some instances, such as root apices, BR and auxin have opposing localization patterns and effects on gene expression (Chaiwanon and Wang 2015). Conversely, in seedlings the two hormones have synergistic effects, where interactions between the BR signaling repressor BIN2 and the auxin signaling repressor ARF2 can alter the seedling’s sensitivity to either hormone (Vert et al. 2008). Additionally, high auxin levels from *yucca* mutants can reduce the effects of BR on hypocotyl elongation by saturating growth responses (Nemhauser et al. 2004). In fact, our live imaging indicates that sepal elongation *drmy1* is more sensitive to BL or BRZ treatments than WT, which has much smaller changes in growth rates and patterns (Figure 6). Perhaps the reduced auxin signaling levels in *drmy1* (Zhu et al. 2020), while not altered by BR treatments (Figure S5C), render *drmy1* more susceptible to both endogenous and exogenous alterations to BR signaling. Additionally, perhaps BL does not alter outer sepal length because *drmy1* outer sepals have higher and more robust auxin levels (Zhu et al. 2020) and elongation is already saturated while inner sepal elongation is not. It stands to reason, then, that inhibiting BR biosynthesis with BRZ would have a much larger impact on the outer sepal because it has more signaling to lose than the inner sepal.

It is also documented that cytokinin signaling is increased in *drmy1* because reduced translation levels lead to low levels of cytokinin signaling components including ARR7 and AHP6, which normally undergo rapid synthesis during sepal initiation (Kong et al. 2024b). It is possible that this same mechanism impacts regulators of BR signaling such as BIN2 to alter BR signaling levels and sensitivities. Recent advances in BR research also suggests that uneven inheritance of BR levels in *Arabidopsis* root cell divisions can increase overall BR activity by circumventing the negative feedback loops typically associated with high levels of BR signaling (Vukašinović et al. 2025). It is possible that the higher variation in BR signaling (Figure 4B) associated with *drmy1* creates a similar mechanism for spatially segregating BR biosynthesis from cells with high BR signaling, thus increasing BR sensitivity. In fact, our spatial RNAseq dataset suggests that the BR biosynthesis gene *DWF1* is expressed most highly not in the regions with initiating sepals themselves, but in the adjacent spots just interior (Figure 3G).

Results from this study reinforce the importance of coordinated growth patterns for proper organ development. It is clear that the proper regulation of multiple hormones such as auxin, cytokinin, and brassinosteroids is crucial for robust growth. This work paves the way for future studies to further investigate the complex dynamics and cross talk between hormones and how they lead to coordinated growth during sepal development.

### Limitations of this Study

We recognize that the untargeted spatial RNAseq technology available when these experiments were performed did not have enough resolution to identify transcriptomic differences between different parts of the same flower bud. Future experiments can address this by using newer untargeted methods with higher resolution such as Visium HD, slide-seq, stereo-seq, etc. or targeted methods such as Xenium, MERFISH, (Chen et al. 2022, Rodriques et al. 2019, Chen et al. 2015, Moffitt et al. 2016; Nagendran et al. 2023). Additionally, many published BR biosynthesis and signaling reporters (such as reporters for DWF4, CPD, BIN2, and BZR1) have little or no expression in flower buds, suggesting the promoters/enhancers guiding the expression patterns are more complex in these tissues. Future experiments will be crucial for identifying the link between *drmy1* and the increase in BR signaling noted here, as well as to understand the mechanism behind the differential responses between the outer and inner sepal to BR signaling levels.

## Acknowledgements

We thank Lanxi Hu, Michelle Heeney, Ruqiang Zhang, Mike Scanlon, Aaron Shipman, Lukas Evans, Si Chen, Bella Burda, Maura Zimmerman, Paul Kuehnert, Kaylee Bagdan, and Avilash Singh Yadav for their comments and feedback on this manuscript. We thank the Arabidopsis Biological Resource Center, Zhiyong Wang, and Sigal Savaldi-Goldstein for seeds. We thank Trevor Tivey and Lukas Evans for their technical expertise and Kevin Cox and Iwijn De Vlaminck for their advice aiding in single cell and spatial transcriptomics. We thank Jan Lammerding for the use of their cryostat. We thank the Genomics Facility (RRID:SCR_021727) and the Imaging Facility (<u>RRID:SCR_021741</u>) of the Biotechnology Resource Center of Cornell Institute of Biotechnology for their help with sequencing and imaging experiments.

## Funding

Research reported in this publication was supported by the National Science Foundation NSF EF-2222434 to AHKR, NSF DGE-2139899, National Institute Of General Medical Sciences (NIGMS) of the National Institutes of Health (NIH) under Award Number R01GM134037 and NIH S10OD032251. The content is solely the responsibility of the authors and does not necessarily represent the official views of the National Institutes of Health or other funders.

## Availability of data and materials

The datasets supporting the conclusions of this article will be available in the National Center for Biotechnology Information Gene Expression Omnibus repository upon publication.

## Materials and Methods

### Plant Material

All genotypes are in the Columbia-0 background unless stated otherwise. The following lines were generated as previously described: *ap1 cal* AP1-GR (Ler background) (Yu et al. 2004; Wellmer et al. 2006) , *drmy1 ap1 cal* AP1-GR (Ler background, *drmy1* introgressed into Ler) (Kong et el. 2024b), *35S::mCitrine-RCI2A* (pLH13) (Robinson et al. 2018), *pUBQ10::mCherry-RCI2A* (Zhu et al. 2020), *DR5::3xVENUS-N7* (Heisler et al. 2005), *pUBQ10::BES1-ypet*/*UBQ10::H2B-mCherry* (Ackerman-Lavert et al. 2021). *bzr1-1D* was previously described by Wang et al. (2002). *det2-1* (CS6159) was obtained from the Arabidopsis Biological Resource Center (ABRC).

### Plant growth

All *Arabidopsis* (*Arabidopsis thaliana*) were grown on Lambert Mix LM-111 soil. Seeds were sown on soil and stratified for about three days at 4°C in darkness before transferring to the growth chambers. *ap1 cal* AP1-GR and *drmy1 ap1 cal* AP1-GR plants were grown in Percival growth chambers at 16°C with constant illumination provided by Philips 800 Series 32 Watt fluorescent bulbs (f32t8/tl841) (∼100 µmol m−2 s−1). All other genotypes were grown in Percival growth chambers at 22°C under long day conditions (16 hr light 8 hr dark) provided by Philips 800 Series 32 Watt fluorescent bulbs (f32t8/tl841) (∼100 µmol m−2 s−1).

### Tissue Collection

Once *ap1 cal* AP1-GR and *drmy1 ap1 cal* AP1-GR plants began to bolt, they were induced with 10 μM Dexamethasone in 0.015% Silwet (vol/vol) three times at roughly 24 hour intervals. About 108 hours (4 ½ days) after the first induction, when flower buds had reached stages 2-3, whole cauliflower-like inflorescences were cut and incubated in protoplasting solution, filtered, pelleted, and washed as described in Satterlee et al. 2020 with the following adjustments: cells were incubated in protoplasting solution for 90 minutes, were passed through a 70 micron filter once and a 40 micron filter twice, and 5 μg/mL final concentration of fluorescein diacetate (FDA) was added after the final resuspension in 50-100 µL wash solution to stain live cells for counting. Two biological replicates were protoplasted per genotype using 0.25-0.6 g of tissue per replicate.

Cells were counted on a Countess 3 FL Automated Cell Counter (Invitrogen) and diluted to allow loading onto the 10× Genomics Chromium Controller using an 8 reaction 3’ HT kit v3 following the manufacturer’s instructions to target ∼10,000 cells per replicate.

For spatial RNAseq with 10x Visium, plants were induced the same way as for scRNAseq. After 4 1/2 days, small pieces of the floral meristem cluster were carefully removed and submerged in Tissue Tek® O.C.T. Compound (Sakura®) in a 10 x 10 x 5 mm cryomold. Air bubbles were carefully removed with a 200 µL pipette tip and 9-12 cluster pieces were arranged on the bottom of the cryomold. O.C.T. embedded tissue was then flash frozen in liquid nitrogen cooled isopentane (2-methylbutane) and stored at -70°C until cryosectioning.

### Single Cell RNAseq Processing

Single cell libraries were sequenced on two P3 flow cells on an Illumina NextSeq 2k 100 bp kits. Raw single cell data reads were converted to fastq format with cellranger v7.0.1 mkfastq and converted to feature matrices with cellranger count using default settings. We used a custom reference by running the TAIR10 v6.0.2 Arabidopsis genome through cellranger mkref using default settings. Matrices were loaded into R version 4.3.0 and analyzed in Seurat v4 (Satija et el. 2015). Data was filtered to only include cells expressing greater than 600 but less than 6000 genes (Figure S1G) and with less than 10% mitochondrial transcripts and 20% chloroplast transcripts. For the remaining cells, PercentageFeatureSet was used to calculate the percentage of transcripts originating from mitochondrial genes (Figure S1H) and chloroplast genes (Figure S1I) for later visualization. Cell cycle was scored using a list of cell cycle genes from Zhang et al. 2021 (Supplementary Data 3) and the CellCycleScoring function in Seurat. To remove the effects of cell cycle on clustering (Figure S1E), the percent variance explained by cell cycle phase for each gene was calculated using the GetVarianceExplained function from the R library “scater” and any gene with >3% variance explained by cell cycle (1,189 genes total) was deleted from the dataset (Supplementary Data 4). Mitochondrial genes were also removed. To check the effects of protoplasting, we used the bulk RNAseq of WT and *drmy1* in the *ap1cal* 35S::AP1-GR background published in Kong et al. (2024b) to check for genes expressed in our single cell dataset that are not expressed in the bulk RNAseq dataset. We identified 2,421 genes in WT and 2,522 genes in *drmy1* potential protoplast induced genes with this method.

The top 5000 variable features were selected with FindVariable Features set to the selection method “vst”. Data was transformed with SCTransform version 2 with 5000 variable features. WT and *drmy1* samples were integrated using FindIntegrationAnchors and with normalization.method set to “SCT” and anchor.features chosen with the command SelectIntegrationFeatures. The anchors were then used with the command IntegrateData set to dims = 1:50 and normalization.method set to “SCT”.

### Dimensionality Reduction

UMAPS were generated in seurat from the integrated scRNAseq data by scaling the data with ScaleData followed by RunPCA set to npcs = 50 and RunUMAP on all 50 principal components. Clustering was performed with FindNeighbors (dims = 1:50) and FindClusters (resolution = 0.4).

### Cluster Annotation

Clusters were annotated by viewing marker gene expression patterns with the built in Seurat functions FeaturePlot and VlnPlot. Marker genes were chosen from several published single cell references (Conde et al. 2022; Zhang et al. 2021) (Supplementary Data 5) and additional marker genes were identified using the “scoreMarkers” function from the scoreMarkers package and setting the mean Area Under the Curve (AUC) threshold to 0.7. (Supplementary Data 6). The most informative markers were then visualized with the dotplot function in Seurat using SCT normalized expression values. Floral meristem atlas mapping was performed as described in Neumann et al. (2022).

### Gene Expression Analysis

Lists of genes that are up and down-regulated in Arabidopsis seedlings in response to exogenously applied hormones were downloaded from Nemhauser et al. (2006).

PercentageFeatureSet was used to calculate the percentage of transcripts in each cell that originate from each list of up or down-regulated genes and plotted with FeaturePlot, VlnPlot, and Dotplot. For dotplots, the dot size was determined by the log base 10 of the inverse of the p-value, with a bonferroni corrected significance threshold of p < 0.000245. Dots with p-values above this threshold were not displayed. Dot color was determined by subtracting the mean value for WT samples from the mean value for *drmy1* samples - thus higher positive numbers represent higher expression levels in WT and higher negative values represent higher expression levels in *drmy1*.

### GO Term Enrichment

We pseudobulked the scRNAseq dataset using the AggregateExpression function in Seurat, grouped by cluster identity and by replicate. Then differential gene expression was performed using the DEseq function in DEseq2 with default parameters. To perform Gene Ontology (GO) Term enrichment on differentially expressed (DE) genes, the DE gene list was first filtered to genes with adjusted p-values less than 0.05, then we imported GO Terms from TAIR “ATH_GO_GOSLIM.txt” into R. We performed GO Term Enrichment for each pseudobulked single cell cluster with the GSEA function (Gene Set Enrichment Analysis) from the clusterProfiler bioconductor package (Wu et al. 2021), setting the maxGSSize to 5000 and the pvalueCutoff to 0.05. The results were visualized with a dotplot in ggplot2, with GO terms and clusters arranged according to similarity using hierarchical clustering via the hclust function in R. A dendrogram for the GO Terms was created using the ggtree function from the ggtree library (Xu et al. 2022).

### Cryosectioning for Spatial RNAseq

Fresh frozen floral meristem clusters stored at -70°C were equilibrated to -10°C in a Thermofischer HM 525 NX cryostat for about 30 minutes. Tissue blocks were cut into 10 micron thick sections and placed on each capture area according to the instructions in the 10x Visium User Guide. Each capture area had a section from a separate block of frozen tissue. For optimization slides, we put 4 *drmy1 ap1 cal AP1-GR* sections and 4 WT *ap1 cal AP1-GR* sections on each slide, while the spatial gene expression slide had 2 sections from each genotype to serve as biological replicates.

### Optimization for Spatial RNAseq

To optimize RNA permeabilization, we used the Visium Spatial Tissue Optimization Slide & Reagent Kit, 4 slides (10x Genomics PN-1000193). We followed the optimization protocol included with the kit with the following adjustments. For all four optimization slides, we skipped the H&E staining steps, instead either not staining tissue (Slide 1) or staining in 0.5 mg/mL toluidine blue in 3.5% ethanol 96.5% water for 30 sec - 4 minutes. Slides were briefly dipped in 1x PBS buffer prepared with RNAse free water to rinse off excess toluidine blue, then extra liquid was allowed to drip off the slide. For slides 2, 3, and 4 we added a pre-permeabilization step according to the instructions in (Giacomello et al. 2017). Briefly, immediately prior to permeabilization we incubate the capture areas in 70 uL of pre-permeabilization mixture (2% Polyvinylpyrrolidone-40, 1x Exonuclease I buffer, 0.19 ug/uL BSA) at 37C for 30 minutes, then rinsed in 100 uL 0.1x SSC buffer. Various permeabilization times between 1 and 30 minutes were tested, with 4-5 minutes determined to be optimal (Figure S9). Brightfield imaging was performed either on a Microbeam laser capture microdissection system (Zeiss) with the 20x objective or a Keyence BZ-X810 with the 20x objective (0.45 NA). Fluorescent cDNA was imaged either on the Keyence BZ-X810 with TRITC filter under the 20x objective (0.45 NA) or on a Zeiss 710 upright confocal microscope with a 20x Plan-Apochromat water-dipping lens (1.0 NA).

### Spatial RNAseq

Visium spatial RNAseq was performed according to the instructions in the user manual with the following notes and alterations. One cryosection slice was placed on each capture area of the Visium slide, each containing approximately 9-12 WT *ap1 cal AP1-GR* or *drmy1 ap1 cal AP1-GR* cauliflower-like clusters. After methanol fixation, H&E staining was replaced with toluidine blue staining as described above, then a coverslip was mounted with a solution of 85% glycerol, 10% RNAse free water, and 0.5% RNAseOUT, followed immediately by brightfield imaging as described above. Pre-permeabilization was performed as described above, and then the user manual was followed for the rest of the protocol using a 5 minute permeabilization time.

### Spatial RNAseq Processing

Paired end libraries were sequenced on one P2 flow cell on an Illumina NextSeq 2k 100 bp kits and one mid-output flowcell on an Illumina NextSeq500 150 bp kit. Reads were converted to fastq format with spaceranger mkfastq and analyzed with spaceranger count v2.0 with default settings. We used a custom reference by running the TAIR10 v6.0.2 Arabidopsis genome through cellranger v7.0.1 mkref using default settings. Brightfield images of capture areas were manually aligned to the capture spots and fiduciary frame in cLoupe browser v6.4.1.

### Spatial Deconvolution

Deconvolution was performed in Seurat v4 by first merging data from all four capture areas, then normalizing with SCTransform (assay = “Spatial”) and creating a PCA reduction with RunPCA. FindTransferAnchors was used to find anchors between the single cell dataset and the Visium dataset, then a predictions assay was generated with the TransferData function using the anchors, single cell cluster identities as the refdata, and the Visium PCA reduction as the weight.reduction, and dims = 1:30. For quality control, we visualized the number of features, UMIs, and percent mitochondrial transcripts on the samples. We found that one of the capture areas with a WT section had abnormally higher mitochondrial transcripts than the others (Figure S4) suggesting cells in this section may have been dying, so we omitted this capture area from downstream analyses.

We used SpatialFeaturePlot to visualize the predicted mapping of each single cell cluster onto the Visium spots. For each Visium spot, we identified which single cell cluster had the highest prediction score and entered this cluster as a new metadata feature and visualized it with SpatialDimPlot. The percentage of transcripts from various hormone up and downregulated genes was calculated for each Visium spot using the same process as for the single cell dataset.

The results were visualized using SpatialFeaturePlot, and in order to generate plots with the same scale for each capture area, we added “& scale_fill_viridis_c(option = "turbo", limits = c(0,upper_limit), oob = scales::squish))” where “upper_limit” is the maximum percentage found among all Visium spots.

### Brassinazole and Brassinolide Experiments

*drmy1* and WT (col-0) plants expressing the pLH13 (*35S::mCitrine-RCI2A*) membrane marker were used right when they began to bolt (bolts 0-3 cm in height). Hormone solutions were created .0025% (v/v) Silwet and either 50 μM brassinazole (50 mM stocks dissolved in DMSO or DMF) (VWR TCB2829-100MG), 400 nM brassinolide (400 μM stock dissolved in DMSO or DMF) (Cayman Chemicals CAYM-21594-25), or 0.1% DMSO or DMF (mock treatments). Plants were dipped in 100 mL baths of their respective treatments’ solution once daily for 1 minute for 6 days. On the 7th day, inflorescences were dissected under a Zeiss Stemi 2000-C Stereo Microscope with a Dumont tweezer (Electron Microscopy Sciences, style 5, no. 72701-D) down to stage 6 or earlier flower buds (Smyth et al. 1990). Dissected inflorescences were placed on 60 mm x 15 mm petri dishes (VWR 25384-092) with live imaging media modified from Hamant et al. (2014). The media was made with 0.5x MS basal salts, 1x Gamborg vitamin mixture, 1% w/v sucrose, 1 mg/mL MES buffer, 1.2% w/v phytoagar (Fisher), and 0.1% Plant Preservative Mixture (Plant Cell Technology) and pH adjusted with 1M KOH to 5.8. For silique images from Figure S5A, 11 days after the first BRZ/BL treatment, the earliest stage 17 siliques and their pedicels were removed from the stem and imaged under a Zeiss Stemi 508 stereomicroscope using a Schott KL 300 LED light source and an Accu-scope camera with CaptaVision+ software.

For single timepoint experiments, plants were imaged immediately on a Zeiss 710 upright confocal microscope with a 20x Plan-Apochromat water-dipping lens (1.0 NA). Due to frequent silencing of the pLH13 membrane marker, plants in single timepoint experiments were stained with a 0.1 mg/mL propidium iodide and 0.1% Silwet solution for 5 minutes and rinsed 3-5 times in DI water before imaging.

For live imaging experiments, the media plates also contained either 5 μM brassinazole, 40 nM brassinolide, or 0.1% v/v DMF (mock). Images were taken on a Leica Stellaris 5 confocal microscope with a 25x Plan-Apochromat water-dipping lens (0.95 NA) every 12 hours for 36 to 60 hours. Between timepoints, samples were placed back in the same long day growth chamber plants were originally grown in (see “Plant Growth”).

The following laser and wavelength were used in confocal imaging. For chlorophyll, excitation 488 nm, emission 647-721 nm. For PI, excitation 514 nm, emission 566-650 nm. mCherry, excitation 594 nm, emission 600–659 nm. For mCitrine/ypet/VENUS/, in *35S::mCitrine-RCI2A*, excitation 514 nm, emission 519-580 nm; in *DR5::3xVENUS-N7* and *pCUC1::3xVenus-N7*, excitation 514 nm, emission 519-569 nm.

### Cycloheximide Treatment

Cycloheximide treatments were performed as described in Kong et al. (2024b).

### Image Analysis

For single timepoint measurements of sepal lengths and outer/inner sepal ratios, confocal image stacks were first converted to maximum intensity projections. These projections were loaded into Fiji v1.53c (Schindelin et al. 2012) and the scale was set using the scale bar included in the projections. The line tool was used to measure the length from the back (proximal side) of the sepal to the longest point of the sepal tip (distal side) for inner and outer sepals. The diameter (length) of the entire bud from the back of the outer sepal to the back of the inner sepal was also measured (Figure 5B). Boxplots were generated in R with ggplot2 and statistics calculated with a Wilcoxon two sided signed rank test.

For live imaging experiments, early stage 3 buds that had just initiated outer sepals were chosen to serve as the 0 hour timepoint for quantifying sepal growth. To help control for sepal initiation timing variability in *drmy1*, buds that had not begun initiating inner sepals at the 12 hour timepoint were excluded from analysis. Confocal stacks were once loaded into ImageJ and converted to .tif files before loading into MorphographX 2 (Barbier de Reuille et al. 2015; Strauss et al. 2022). Images were brightened once or twice with Stack/Filter/Brighten Darken (amount = 2) and blurred with Stack/Filters/Gaussian Blur Stack (x = y = z = 1) 2-3 times.

Stack/Lyon/Init level set (up threshold = down threshold = 2) was run followed by Stack/Lyon/Level Set Evolve (alpha = 0, beta = 0.1, lamda = 10, dt = 100, epsilon = 1.5, threshold = 0.002, max steps = 5, view steps = 5) 2-3 times. Stack/Morphology/Edge Detect Angle (fill holes = no) was then applied, and Stack/Morphology/Closing (x = y = z = 3, Round = no) used to close holes. Any material outside of the bud of interest was manually removed using the voxel edit tool. A mesh was created using Mesh/Creation/Marching Cubes Surface (cube size = 5, threshold = 20000), then refined using Mesh/Structure/Smooth Mesh (passes = 10, walls only = no) and Mesh/Structure/Subdivide functions three times each. Cell outlines were projected onto the mesh using Mesh/Signal/Project Signal (use absolute = no, min signal = 0, max signal = 60000) with a variable depth optimized to capture the best signal from epidermal cells from each bud - most often min distance = 1 μm and max distance = 3 μm was ideal. For segmentations of every cell, Mesh/Signal/Gaussian Blur (radius = 0.5 μm) was used to blur the cell membrane signal slightly, then cells were manually seeded, then segmented with Mesh/Segmentation/Watershed Segmentation (steps = 50000) and Mesh/Segmentation/Fix Corner Triangles. For analyses where entire sepals growth rates were analyzed, each sepal was segmented as if it were one big cell. This was achieved by manually defining the boundaries of the sepals at the latest (36 hour) timepoints following existing cell boundaries. Then earlier timepoints were segmented by following the same cell boundaries to create the sepal boundaries.

To calculate growth rates between timepoints, a previous timepoint would be loaded as a second mesh and parent labels manually set by aligning the two buds and using the “grab labels from other surface” tool. Errors in parent tracking and segmentation were identified using Mesh/Cell Axis/PDG/Check Correspondence and manually fixed by deleting labels and re-segmenting as needed. Heat maps were made using Heat Map Classic with type set to Area, visualization to geometry, “change map” option checked with “ratio” and “decreasing” chosen in the drop down menus and “use manual range” set to minimum 0 maximum 2.5. For whole sepal analyses, the growth rate heatmaps were exported using Mesh/Attributes/Save to CSV extended. Absolute growth amounts were calculated by copying the area of each sepal from the previous timepoint into the CSV file and subtracting the current timepoint’s area from the previous area for each sepal. Percentage contributions to total sepal growth were calculated by summing the absolute growth amounts from all sepals, then dividing each sepal’s absolute growth by the summed growth. Sepal ratios were calculated by simply dividing the area of the outer sepal by the area of each other individual sepal. However, in cases where there were two inner sepals instead of one, both inner sepals together were treated as one sepal to prevent inflating the outer/inner sepal ratio.

For BES1/H2B ratiometric images, confocal stacks were once again converted to .tif files to load into MorphoGraphX. The H2B channel was loaded first, and the main stack was copied to the work stack and signal outside of the bud(s) of interest was removed with the voxel edit tool. Stack/Filters/Gaussian Blur Stack ( x=y=z radius = 0.8 μm) was applied, then seeds created with Stack/Segmentation/Detect Local Maxima (x = y = z radius = 0.5, start label = 2, threshold = 3000, value = 60000). Meshes were created with Mesh/Creation/Mesh From Local Maxima (radius = 2 μm). Many overlapping nuclei meshes are inadvertently created with this method, in which case one overlapping nucleus is chosen at random and manually removed by removing the label, using Mesh/Selection/Select Unlabeled, and hitting backspace to delete. Labels were added to each bud corresponding to their developmental stage by using the lasso tool to select all of the nuclei in a bud and using Mesh/Lineage Tracking/Set Parent to assign a unique numeric parent label based on the bud’s stage. Mesh/Signal/Project Signal (min = 0 μm, max = 2 μm, min signal = 0, max signal = 100000, use absolute = no) projected the nuclear H2B signal onto the meshes, which was then quantified with Mesh/Heat map/Measures/Signal/Signal Total. Then the BES1 channel was loaded into MorphographX and projected onto the mesh in the same way, but quantified using Mesh/Heat map/Measures/Signal/Signal Interior. This quantification is functionally the same as Signal Total in this case, but allows us to store the BES1 signal as a separate attribute, at which point both can be exported together with Mesh/Attributes/Save to CSV extended. The BES1/H2B signal ratio was calculated for each ratio by dividing the Signal Interior column (BES1) by the Signal Total column (H2B). The ratio can be visualized by using Mesh/Heat map/Heat Map Load and choosing the ratio column, and using Mesh/Heat map/Heat Map Set Range (min = 0 max = 4) to make the scale consistent between images.

Sepal initiation timing was performed as detailed in Kong et al. (2024b).

## Notes

### Competing Interest Statement

The authors have declared no competing interest.

