## Supplementary Figures for "Brassinosteroids mediate proper coordination of sepal elongation"

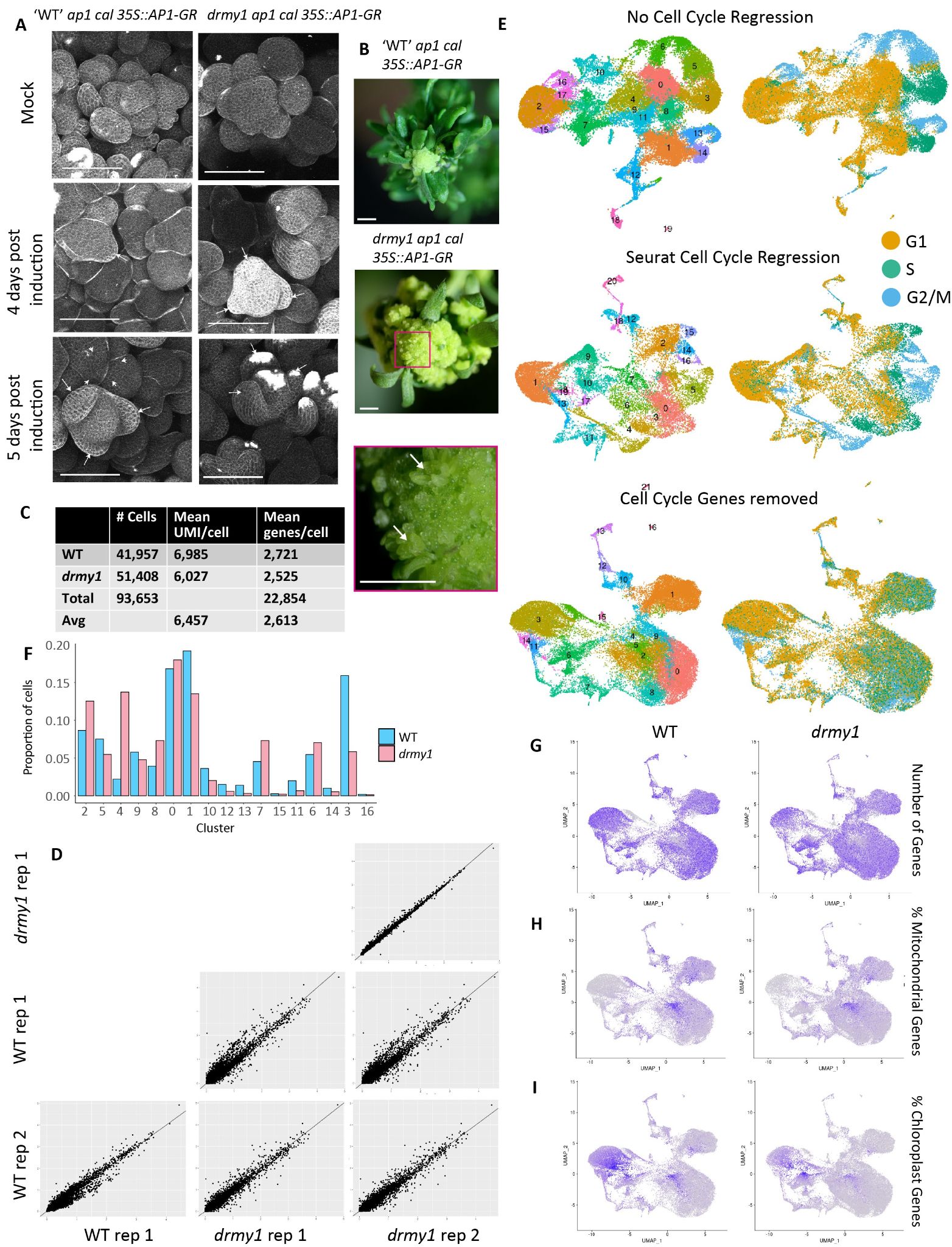

Figure S1: Single cell RNAseq quality control

A) Confocal images of propidium iodide stained WT and *drmy1* floral meristem clusters in the *ap1 cal* 35S::AP1-GR background 4 days and 5 days after induction with 10 μM dexamethasone or mock. Examples of initiating sepals marked with arrows.

B) Stereomicroscope images of pre-induction WT and *drmy1* floral meristem clusters in the *ap1 cal* 35S::AP1-GR background. Note the larger numbers of leafy bracts present in WT, also note the older “escaped” flowers in *drmy1* that began initiation before dex induction marked by arrows.

C) Table of cell counts, mean unique molecular identifiers (UMIs), and mean number of genes detected per cell.

D) Correlation between the average normalized expression of each gene in WT and *drmy1* replicates.

E) UMAPs showing the clustering (left column) and cell phase (right column) of each cell, demonstrating the effects of cell cycle on clustering and how different methods of removing cell cycle effects (no regression on top, Seurat cell cycle regression using the “vars.to.regress” argument in the ScaleData function in the middle, and complete removal of genes with > 3% variance explained by cell cycle phase on the bottom).

F) Proportion of cells from each cluster in WT and *drmy1* single cell data.

G- I) Feature plots of various quality control metrics including the total number of genes detected (G), the percentage of transcripts from mitochondrial genes (H), and chloroplast genes (I) in each cell. Note the relatively high proportion of chloroplast genes in cluster 3, which helped identify this as likely mesophyll.

Scale bars 100 μm (A), 1 mm (B)

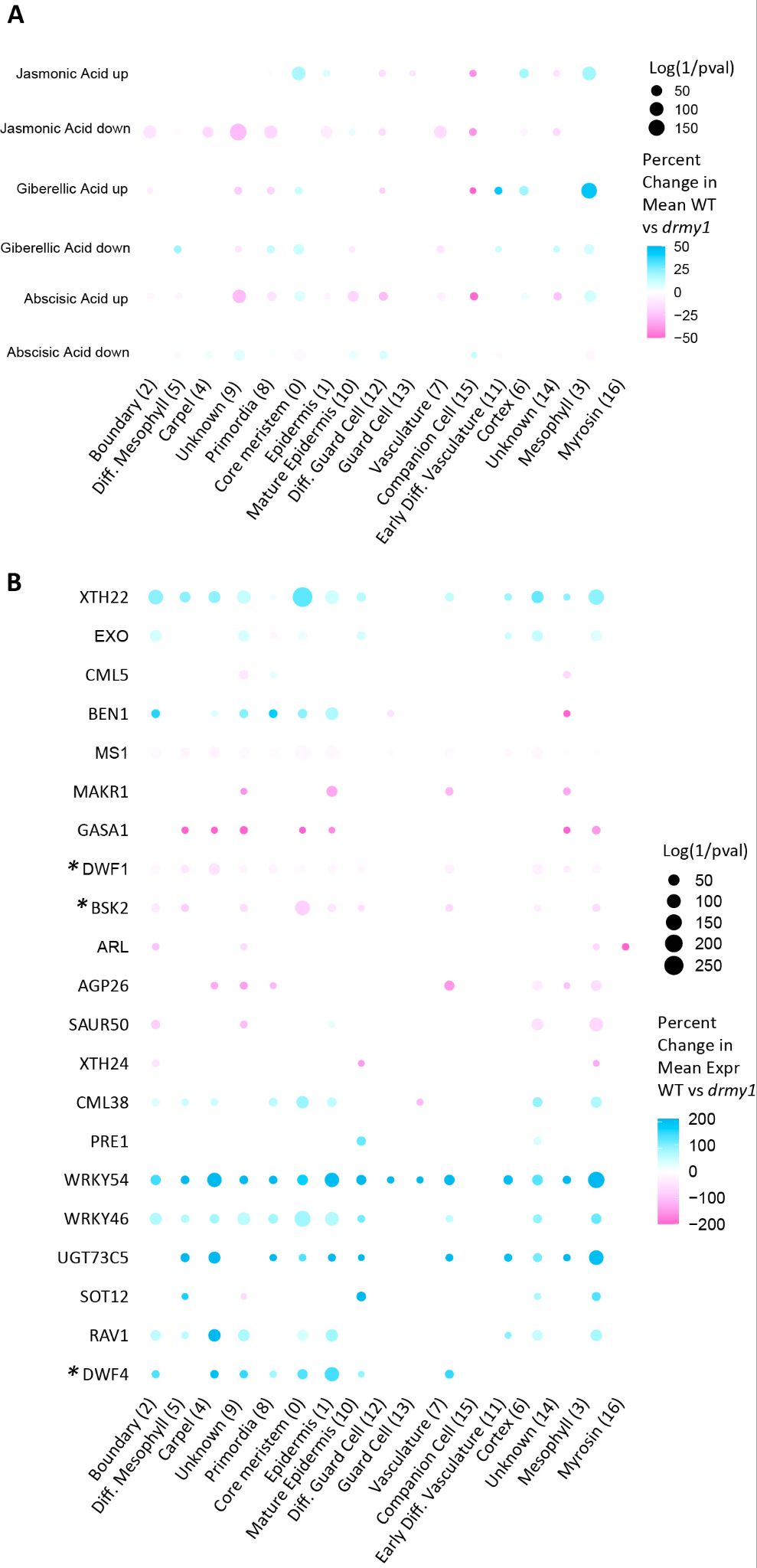

Figure S2: Hormones up and downregulated in *drmy1*.

A) Dotplot showing the differences in expression of genes up and down-regulated by jasmonic acid, Abscisic acid, and gibberellic acid. For each cell, the percentage of transcripts that come from the set of genes up or downregulated by applications of these hormones according to Nemhauser et al. (2006) is calculated, then the mean of all the cells in each cluster is calculated for WT and *drmy1*. The size of each dot represents the log of the inverse of the p-value between the mean of the two genotypes with a Bonferroni adjusted threshold of p < 0.000245. The color of each dot represents the percentage increase or decrease of the mean of *drmy1* compared to WT.

B) Dotplot showing the differences in expression of genes differentially expressed in *drmy1* with gene ontology terms that mention brassinosteroids. The size of each dot represents the log of the inverse of the p-value between the mean expression of the two genotypes with a Bonferroni adjusted threshold of p < 0.00014. The color of each dot represents the percentage increase or decrease of the mean of *drmy1* compared to WT. Stars indicate genes in the brassinosteroid biosynthesis or signaling pathways, which includes *DWF1* (biosynthesis), *DWF4* (Biosynthesis, negatively regulated by brassinosteroids), and *BSK2* (signaling).

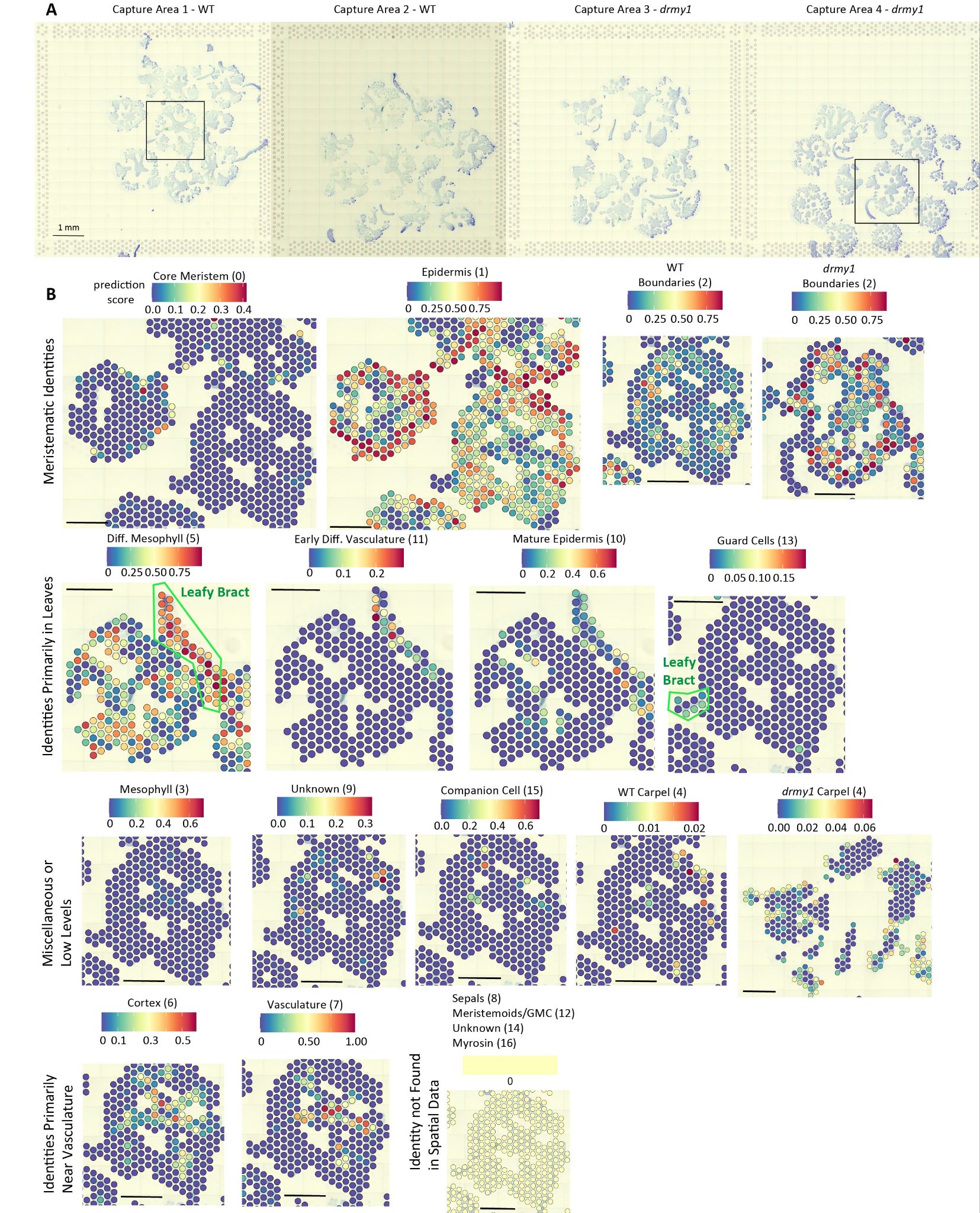

Figure S3: Cell identity patterns in Visium spatial RNAseq from spatial deconvolution.

A) Brightfield images of all four Visium capture areas. Boxes show the representative areas used for visualizing trends in Figure 3. Scale bar 1 mm.

B) Prediction scores of how well each single cell cluster maps to each Visium spot. Organized by similar expression patterns. Scale bars 500 μm.

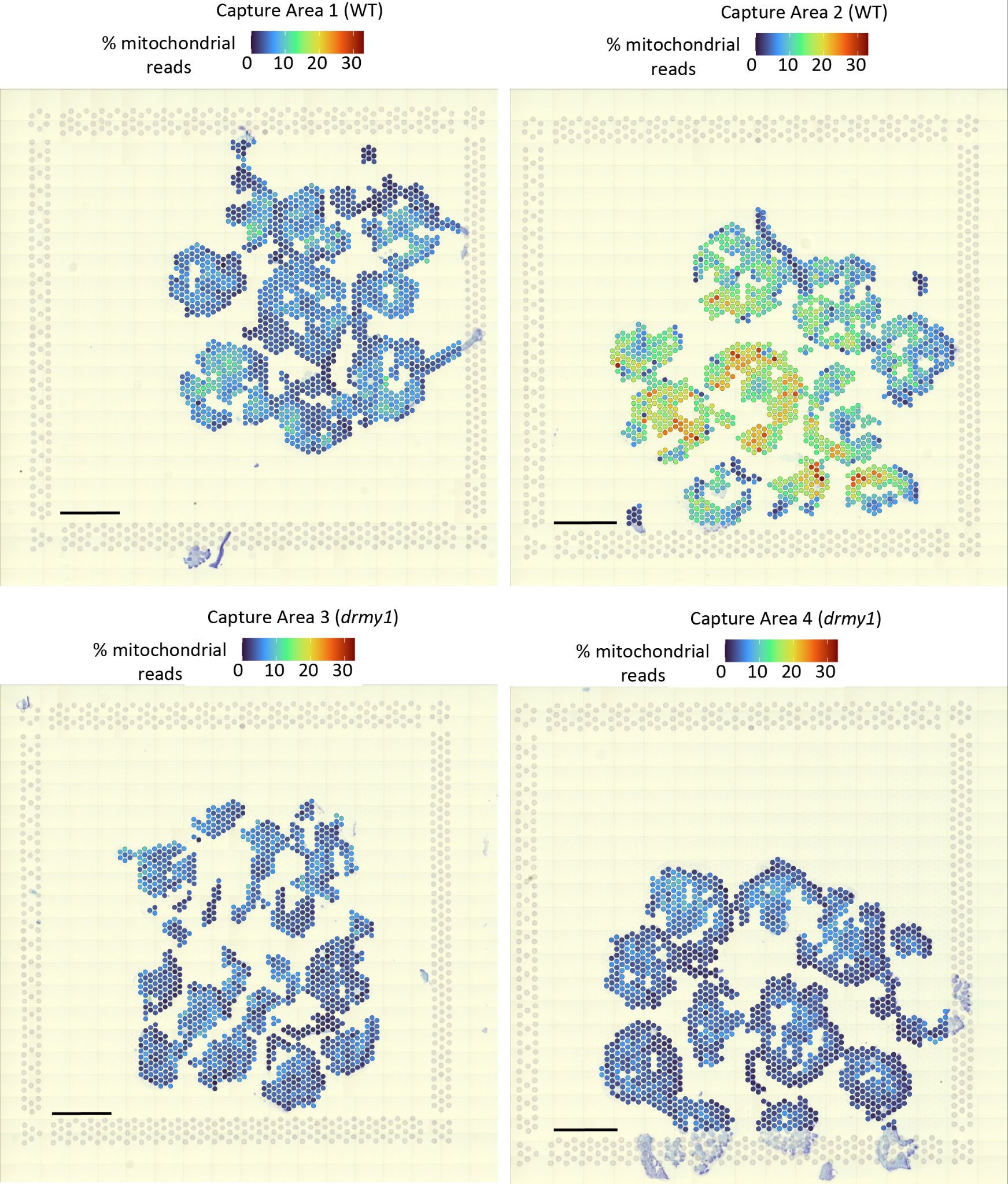

Figure S4: WT capture area 2 has high numbers of mitochondrial transcripts

Percentage of transcripts coming from mitochondrial genes. Note how capture area 2 (a WT replicate) has much higher overall mitochondrial read percentages.

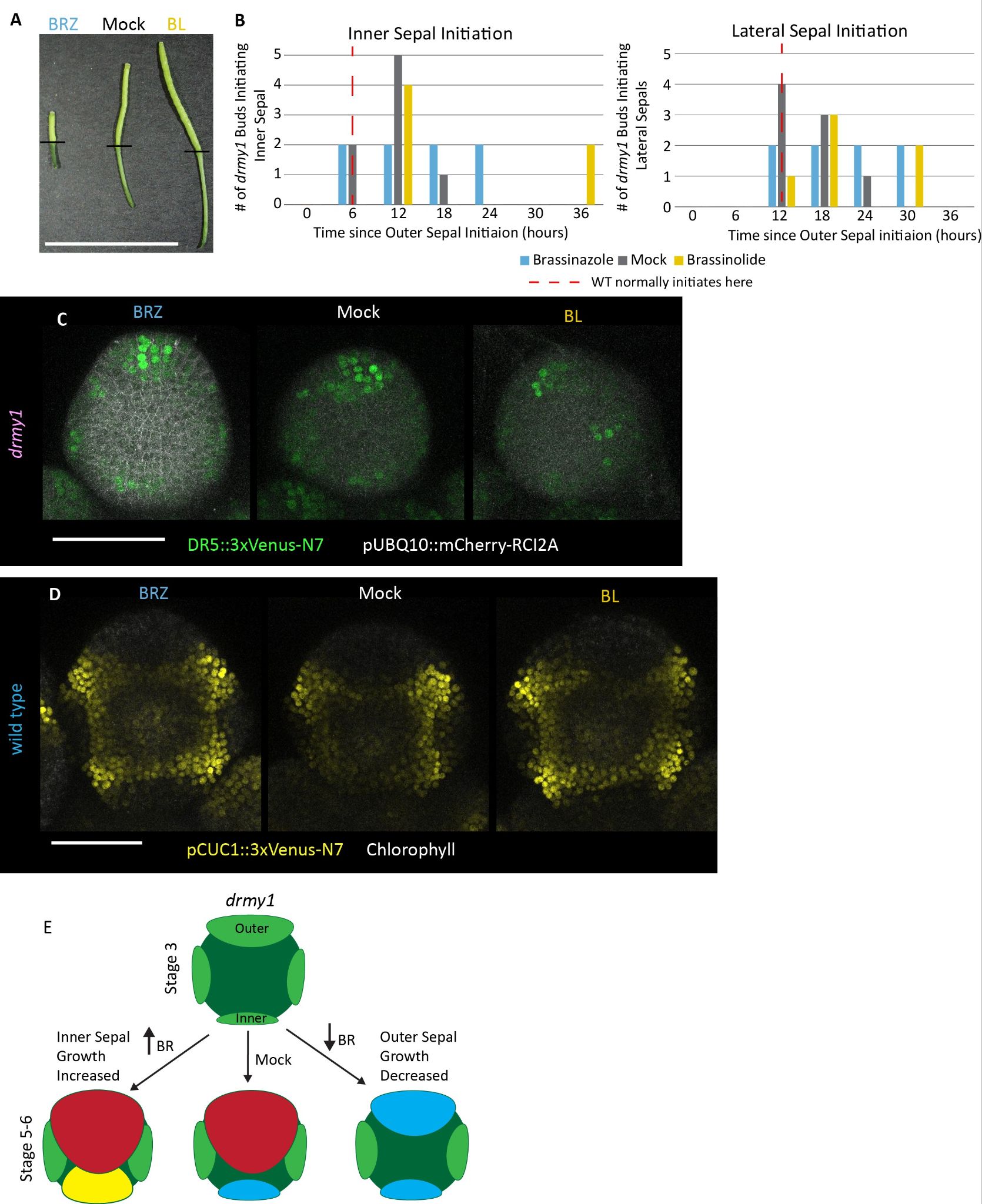

Figure S5: Altering BR levels does not rescue sepal initiation timing or auxin signaling in *drmy1* and does not alter CUC1 expression in young flower buds.

A) Early stage 17 siliques just after other floral organs have fallen off the silique 11 days after treatment with 50 μM brassinazole (BRZ), 400 nM brassinolide (BL), or mock. Black lines indicate the pedicel/silique junction. Note both pedicels and silique lengths elongated by BL treatment and shortened by BRZ treatment, as expected.

B) Timing of initiation of the inner and lateral sepals relative to the initiation of the outer sepal in BRZ, BL, and mock treated *drmy1* buds. In WT, inner sepals would normally initiate 6 hours after the outer sepal, and laterals 12 hours after outer sepal (denoted by red dashed lines).

C) *DR5::3xVenus-N7* expression in stage 2 *drmy1* flower buds under BRZ, BL, and mock treatments.

D) *pCUC1::3xVenus-N7* expression in stage 3 WT flower buds under BRZ, BL, and mock treatments.

E) Model describing the effect of increasing or decreasing BR signaling on the growth of the inner and outer sepals of *drmy1.*

Scale bars 1 cm (A) or 50 μm (C) and (D).

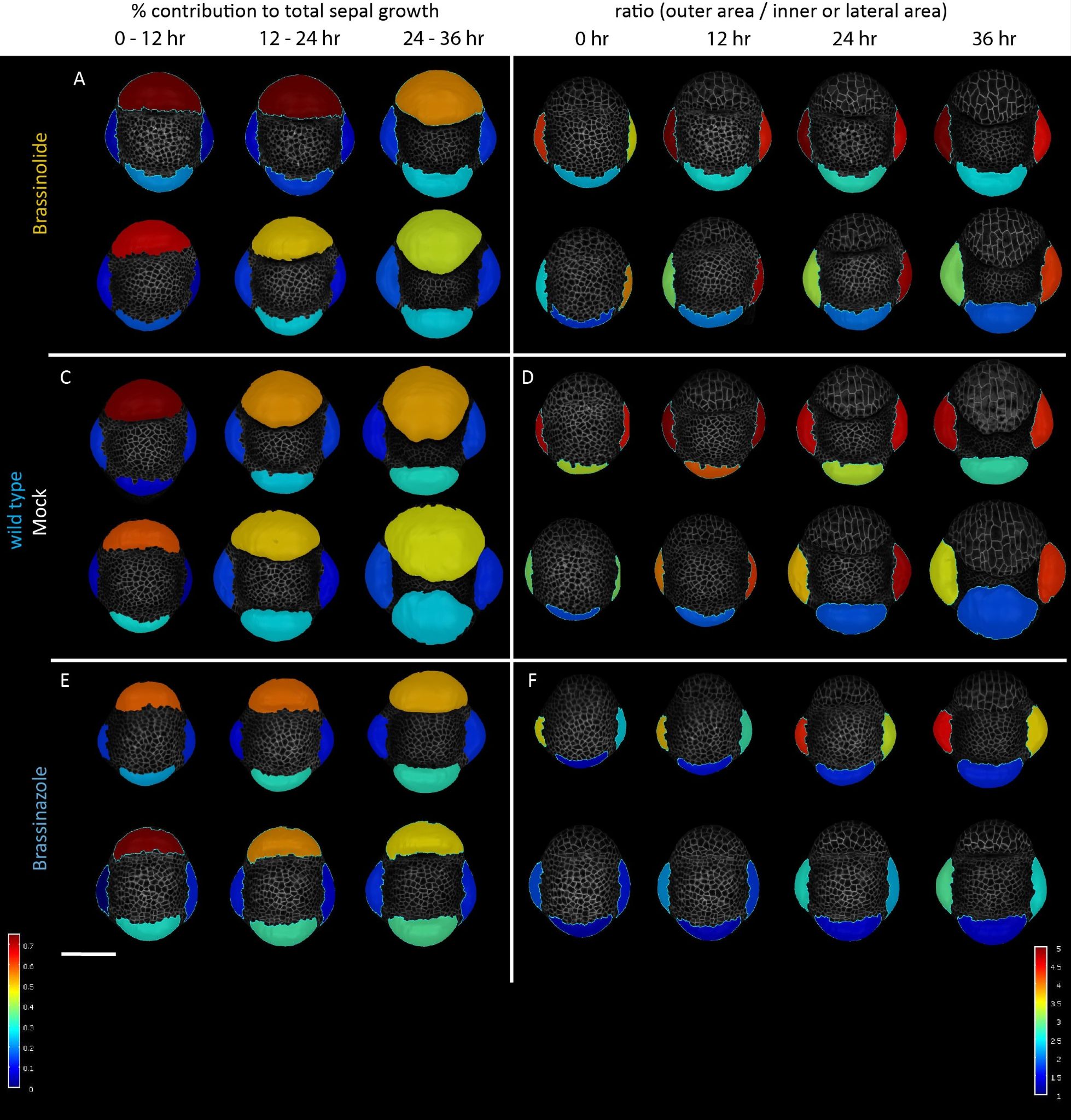

Figure S6: Live imaging replicates of wild type flower buds.

A, C and E) Proportion of total sepal growth contributed by each sepal in WT bud replicates treated with 400 nM brassinolide (BL) (A), mock (C), or 50 μM brassinazole (BRZ) (E) at 12 hour time intervals.

B, D, and F) Corresponding ratio between the area of the outer sepal to the area of the inner and each lateral sepal under BL (B), mock (D), and BRZ (F) treatments in WT replicates.

Scale bars 50 μm.

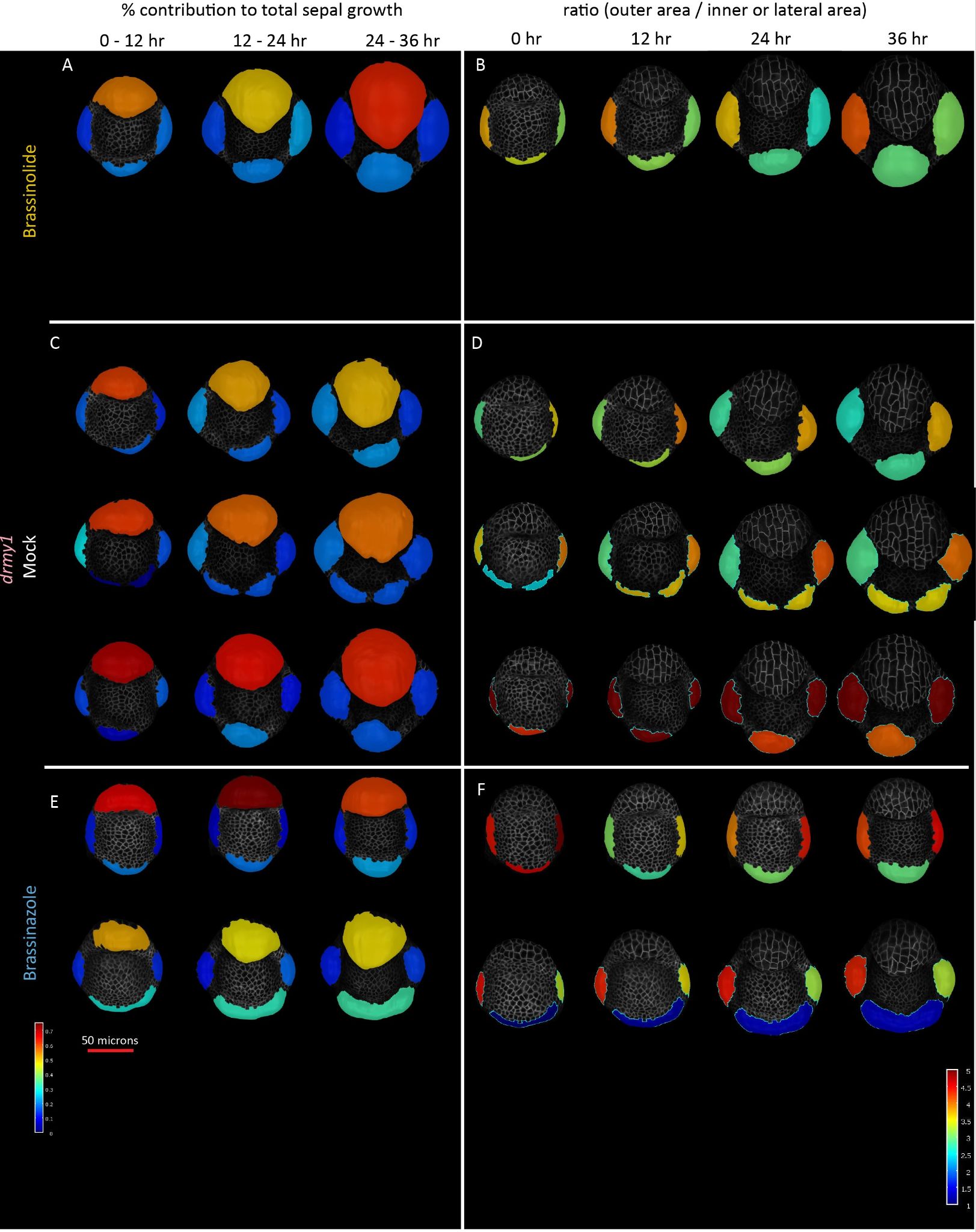

Figure S7: Live imaging replicates of *drmy1* flower buds.

A, C and E) Proportion of total sepal growth contributed by each sepal in *drmy1* bud replicates treated with 400 nM brassinolide (BL) (A), mock (C), or 50 μM brassinazole (BRZ) (E) at 12 hour time intervals.

B, D, and F) Corresponding ratio between the area of the outer sepal to the area of the inner and each lateral sepal under BL (B), mock (D), and BRZ (F) treatments in *drmy1* replicates. Note the correlation between the proportion of sepal growth coming from the outer sepal and the magnitude of the corresponding outer/inner sepal ratio.

Scale bars 50 μm.

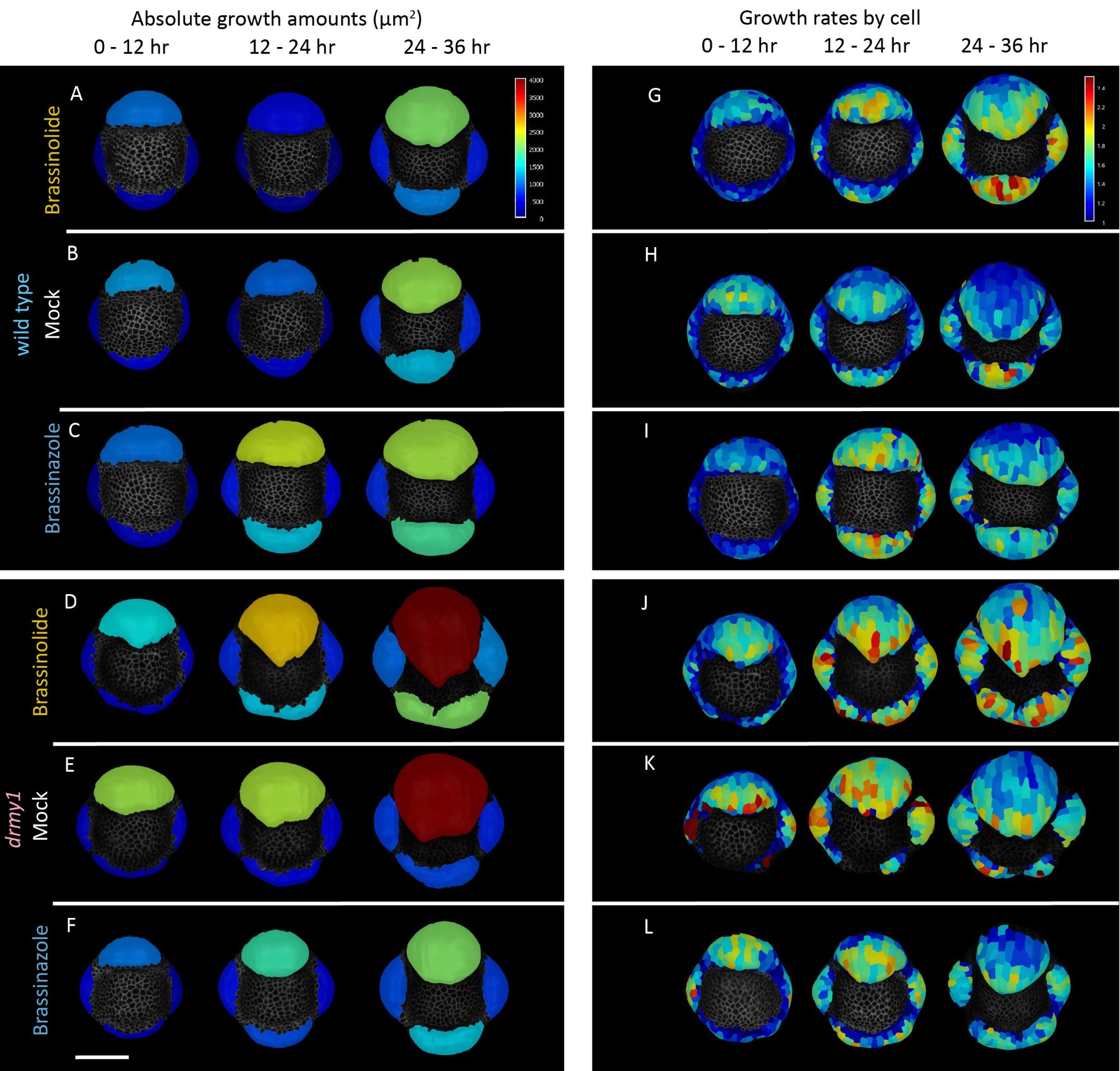

Figure S8: Absolute growth of sepals and growth rates of individual cells confirms that outer sepals overgrowth is diminished by brassinazole, inner sepal growth increased by brassinolide.

A-C) Absolute growth amounts during each 12 hour interval in WT buds treated with 400 nM brassinolide (BL) (A), mock (B), or 50 μM brassinazole (BRZ) (C).

D-F) Absolute growth amounts during each 12 hour interval in *drmy1* buds treated with BL (D), mock (E), or BRZ (F).

G-I) Growth rates of each cell during each 12 hour interval in WT buds treated with BL (G), mock (H), or BRZ (I). Note: these samples do not all correspond to the same samples in the A-C.

J-L) Growth rates of each cell during each 12 hour interval in *drmy1* buds treated with BL (J), mock (K), or BRZ (L). Note: these samples do not all correspond to the same samples in the D-F.

Scale bars 50 μm

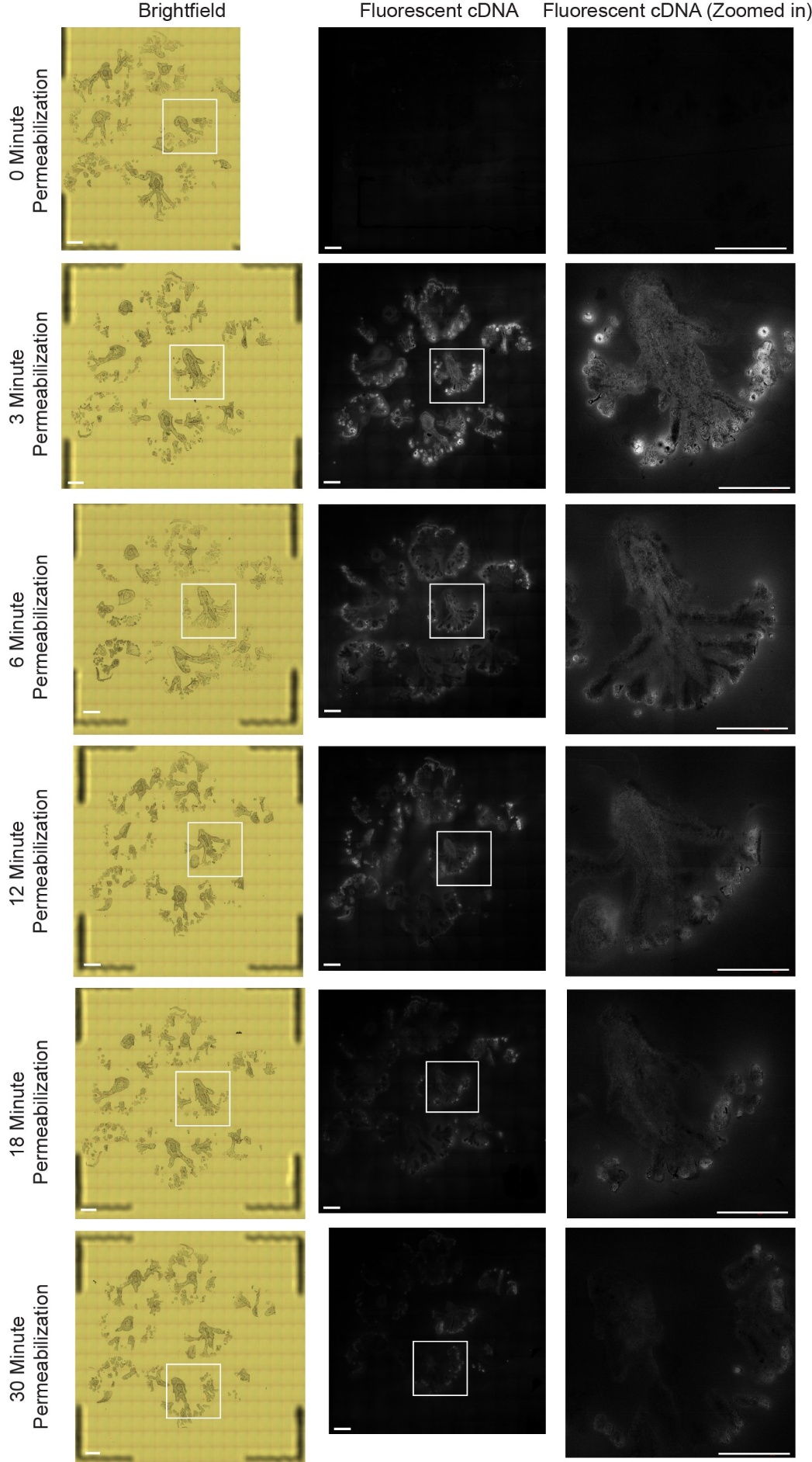

**Figure S9: Optimization of permeabilization times indicates 3-5 minutes yields the best RNA signal**. Stitched brightfield and confocal images of the capture areas and of selected areas. More fluorescent cDNA signal represents better RNA release from permeabilization.

Scale bars 500 μm
